# Systematic assessment of the accuracy of subunit counting in biomolecular complexes using automated single molecule brightness analysis

**DOI:** 10.1101/2021.11.22.469513

**Authors:** John S H Danial, Yuri Quintana, Uris Ros, Raed Shalaby, Eleonora G Margheritis, Sabrina Chumpen Ramirez, Christian Ungermann, Ana J Garcia-Saez, Katia Cosentino

**Author notes:** To whom correspondence should be addressed: Katia Cosentino, John S H Danial and Ana J Garcia-Saez. These authors contributed equally (John S H Danial, Yuri Quintana).

## Abstract

Analysis of single molecule brightness allows subunit counting of high-order oligomeric biomolecular complexes. Although the theory behind the method has been extensively assessed, systematic analysis of the experimental conditions required to accurately quantify the stoichiometry of biological complexes remains challenging. In this work, we develop a high-throughput, automated computational pipeline for single molecule brightness analysis that requires minimal human input. We use this strategy to systematically quantify the accuracy of counting under a wide range of experimental conditions in simulated ground-truth data and then validate its use on experimentally obtained data. Our approach defines a set of conditions under which subunit counting by brightness analysis is designed to work optimally and helps establishing the experimental limits in quantifying the number of subunits in a complex of interest. Finally, we combine these features into a powerful, yet simple, software that can be easily used for the stoichiometry analysis of such complexes.

## INTRODUCTION

Assembly into nanoscopic oligomeric complexes is a common mechanism that allows biomolecules to perform their cellular activities^1–3^. Determining the structural organization of these complexes is paramount to understanding their functions. High resolution structural characterization methods, such as X-Ray diffraction or Cryo Electron microscopy, provide angstrom-resolution atomic maps but require the biological complex under study to be purifiable with a high yield, to preserve its entirety and, possibly, its native physiological environment during the purification step, and to be stoichiometrically homologous. Super-resolution microscopy methods can chart the architecture of many of these nanoscopic complexes directly inside their cellular environments with nanometer resolution ^4–7^, however, precise molecular counting using super resolution microscopies is a formidable task that remains challenging ^5,8–12^.

Single molecule fluorescence analysis has emerged as a powerful strategy to measure the stoichiometry of small and large biomolecular complexes^13–20^. Two major approaches comprising subunit counting by this analytical toolkit are known as stepwise photobleaching^21^ and single molecule brightness analysis^22^. In stepwise photobleaching analysis, the number of photobleaching steps exhibited by a single oligomer are counted and correlated with the number of subunits contained within. Counting the number of photobleaching steps has, traditionally, been performed manually or by the use of some algorithms^23^. However, both approaches require trained users able to isolate actual photobleaching steps from artefacts derived from high noise levels, presence of one or several photo blinking steps and temporal variations in the intensity of the excitation source. These problems were recently addressed by training and deploying Convolutional and Long-short-term memory Deep learning Neural Network (CLDNN) to classify different oligomeric species based on the number of photobleaching steps they exhibit^24^. Despite the high accuracy of this network in discerning oligomers with up to 5 sub-units, automated and manual classification based on step counting remains still extremely challenging for larger assemblies.

Single molecule brightness analysis is not limited by the mentioned factors and has the potential to quantify the stoichiometry of small to medium-sized macromolecular complexes^22^. In this method, the number of underlying subunits of an oligomeric species is obtained by comparing its brightness to a calibration curve theoretically calculated from the measured average brightness of monomers. Monomers, selected based on the stepwise photobleaching analysis, can be obtained by different strategies: from a sample with a mixture of different oligomers, from partial bleaching of protein complexes^15^ or from non-activated or mutant forms of the protein of interest, which are unable to oligomerize^17^. Paramount to the accurate quantification of stoichiometry is the selection of ‘clean’ intensity traces of the monomeric species, which are not affected by any intensity variations other than imaging noise. Equally important is the accurate measurement of the brightness of oligomeric species, which is irrespective of high noise, early photo bleaching, presence of multiple photo blinking steps and other nonspecific intensity variations.

Despite the enormous power of single molecule brightness analysis and the unique niche it occupies within the family of methods used to quantity absolute molecular copy numbers, it suffers from a number of important limitations:

1. The fluorescence intensity of the monomeric and oligomeric particles may take a wide range of values. Detecting these particles with high fidelity (i.e. low false negative and positive rates) requires subjective changes of the detection parameters by the end-user. This process hampers the automation of data processing and the accuracy of the eventual subunit counting.
2. The maximum number of resolved oligomers is strictly connected to the quality of the monomer calibration. Therefore, the selection of clean, single-step intensity traces for monomer calibration is paramount to resolve higher order oligomers, however traditionally this step is performed manually. Besides the potential introduction of human errors during the classification process, this is a complicated task due to the need for tens to hundreds of such traces for appropriate calibration, which needs to be repeated for each data set due to any subtle change in the experimental setup or in the sample preparation.
3. Although the theory behind single molecule brightness analysis was extensively scrutinized, the experimental conditions under which this method is designed to operate optimally have never been systematically navigated. This prevented the optimized application of this method and in some cases may have led to incorrect conclusions on the underlying biological system.
4. The accuracy of this method in quantifying multiple stoichiometric occurrences (e.g. dimeric, trimeric, tetrameric etc.) of protein complexes, as well as any change in the proportion of oligomeric species as a function of protein concentration, was not systematically assessed before. This prevented the end-users from understanding the analytical limits of this approach and judge its applicability to the system of their interest.
5. Fitting the intensity distributions to multiple Gaussians may not perfectly match the real data introducing false stoichiometry assessments.

To address these important limitations, we have developed SAS (Stoichiometry Analysis Software), a fully automated software pipeline to analyse the stoichiometry of oligomeric complexes imaged by fluorescence microscopy. By employing SAS for quantifying the number of complex subunits by brightness analysis, we could carefully assess the accuracy of this method and provide the users with guidelines for the optimal experimental and analytical conditions to employ for reliable and accurate stoichiometry measurements of protein complexes.

## RESULTS AND DISCUSSION

### 2.1 Architecture of SAS (Stoichiometry Analysis Software)

SAS uses a simple, but robust, parameter-free, single-molecule particle detection algorithm based on multi- layer convolutional neural network (DeepSinse), which was previously found to exhibit a 4 to 5 times lower false positive and negative rates compared to the best-in-class, domain specific detection algorithm based on wavelet filtering on a remarkably wide range of Signal-to-Noise Ratios (SNRs)^25^. Importantly, SAS requires no human input for optimized detection (**Figures 1a-c, Supplementary Figure 1**, see **methods**). The time-dependent intensity traces of the detected molecules are extracted from the acquired frame stacks by measuring the background-corrected intensities in Regions Of Interest (ROIs) centred around the centroids of each detected particle (**Figure 1c** and **d**, see **methods**). In SAS, data to be processed need to be classified either as “calibration” or “unknown” (**Figure 1b-d**). The “calibration” data set will be used to find the brightness values of monomeric species. To this aim, the extracted intensity traces are fed into a trace annotator that selects clean, single-step traces by calculating and normalizing the gradient (i.e. slope) of each trace and picking up traces with a single peak gradient above a pre-set threshold (**Figure 1e, Supplementary Figure 2**, see **methods**). The selected traces are then used to construct a distribution curve from a kernel density function and fitting it to a Gaussian Mixture Model (GMM) to account for the fact that some of the selected single-step traces are not monomeric due to the photobleaching of two, or more, fluorophores at the same time within the same complex (**Figure 1f**, see **methods**). The mean and standard deviation of the intensity values in the fitted Gaussian curve corresponding to the monomeric population is, subsequently, used to construct an idealized Gaussian mixture that represents the distribution of the higher-order oligomeric species. By over-imposing these multiple Gaussians on the intensity distribution of all detected particles from the “unknown” data set, the proportion of each species is calculated from the area of each Gaussian curve (**Figure 1g**, see **methods**). Finally, the calculated proportions are corrected for incomplete labelling using a binomial probability density function to yield the true stoichiometry of the underlying biological complex (**Figure 1h**, see **methods**). The quality of the monomeric calibration Gaussian curve is critical in the brightness analysis approach as the width of the intensity distribution defines the maximum number of species that can be resolved. The selection of monomeric traces by SAS is reliable even for wide and complex simulated and experimental intensity distributions (see for example the calibration distribution of experimental data in Fig. S3). However, this step needs particular attention and scrutiny by the final end-user.

**Figure 1.**
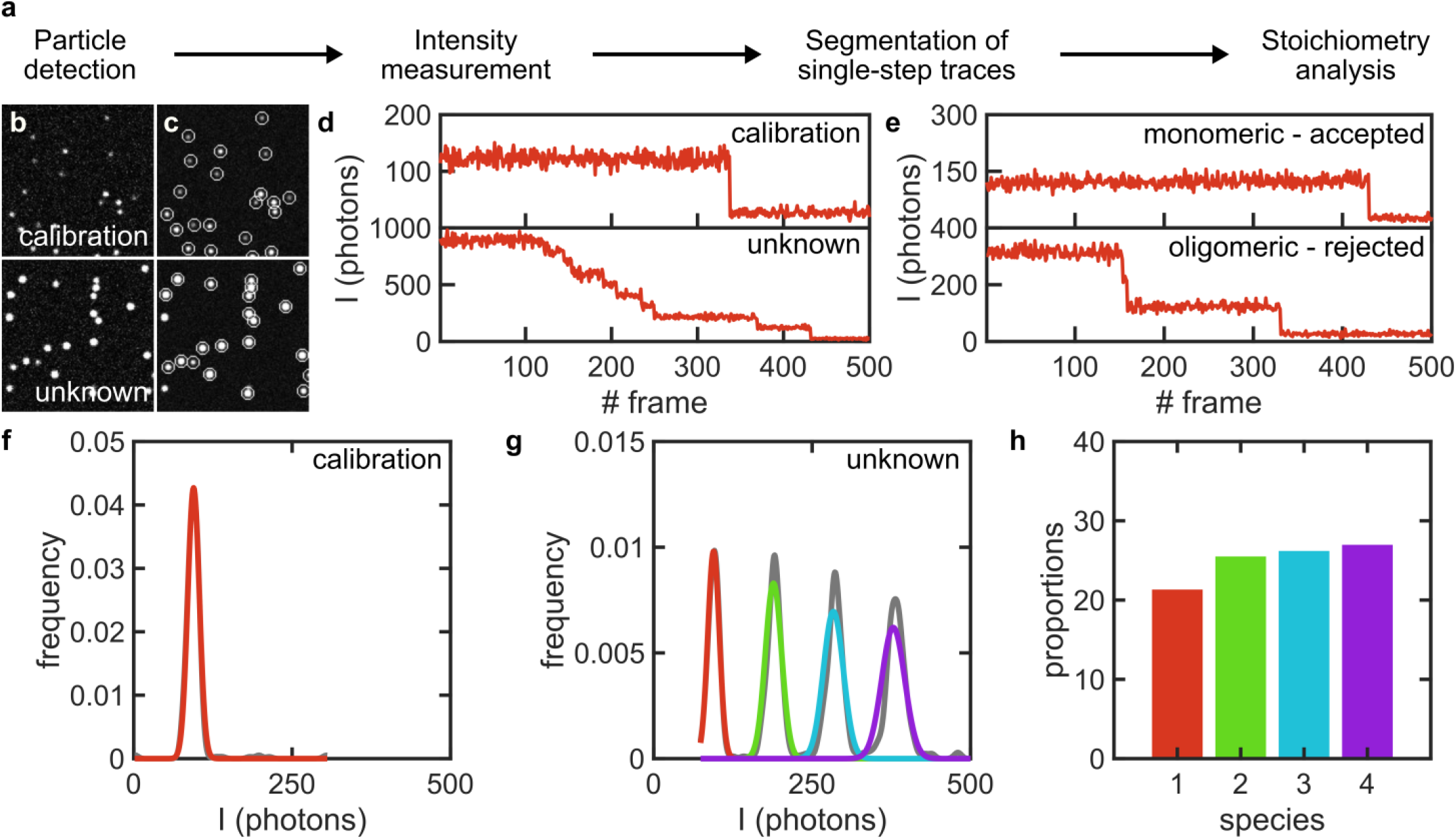
Overview of the mode of operation of SAS. **(a)** SAS workflow. **(b)** Exemplary simulated, ground-truth images of single molecules for a set of calibration (stoichiometry: 50% monomers, 25% dimers and 25% trimers) and unknown (stoichiometry: equal proportions of monomers to 16-mer) species before detection and **(c)** after detection where detected particles are encircled with white circles. **(d)** Exemplary intensity traces of 2 randomly chosen particles from the calibration and unknown datasets after conversion from signal counts to photons. **(e)** Examples of monomeric and oligomeric traces extracted from the calibration dataset and which are automatically annotated by SAS. **(f)** Kernel density function of the intensity distribution underlying the calibration dataset (shown in grey) and the Gaussian curve representing the monomeric population (shown in red). **(g)** Kernel density function of the intensity distribution underlying the unknown dataset (shown in grey) and the Gaussian mixture representing the monomeric population (shown in red, green, cyan and purple). **(h)** Bar graph of the proportion of the species underlying the unknown dataset (color code as in g).

### 2.2 Performance of SAS and assessment of optimal experimental conditions for stoichiometry counting

We then assessed the performance of the software and the accuracy of counting using single molecule brightness analysis by simulating ground-truth data in a wide range of experimental conditions (see **methods**). To this end, we evaluated the error in the calculated *versus* simulated proportions of species when varying the density of particles, SNR, number of subunits per complex (at constant particle density), number of monomeric particles selected for calibration, variation in the intensity of each single molecule, the bin size of the kernel probability distribution function (pdf), the particle intensity distribution width (i.e. sigma) to ROI radius (SRR) from which the intensity traces are extracted and the pixel size (**Figures 2a-h**). Under all simulated conditions, the error in the assignment of oligomeric species did not exceed 15% whilst reaching, in many cases, to lower than 5%. Variations in the density of particles, number of subunits per complex and pixel size did not result in substantial changes to the error (< 3%) within the simulated ranges.

**Figure 2.**
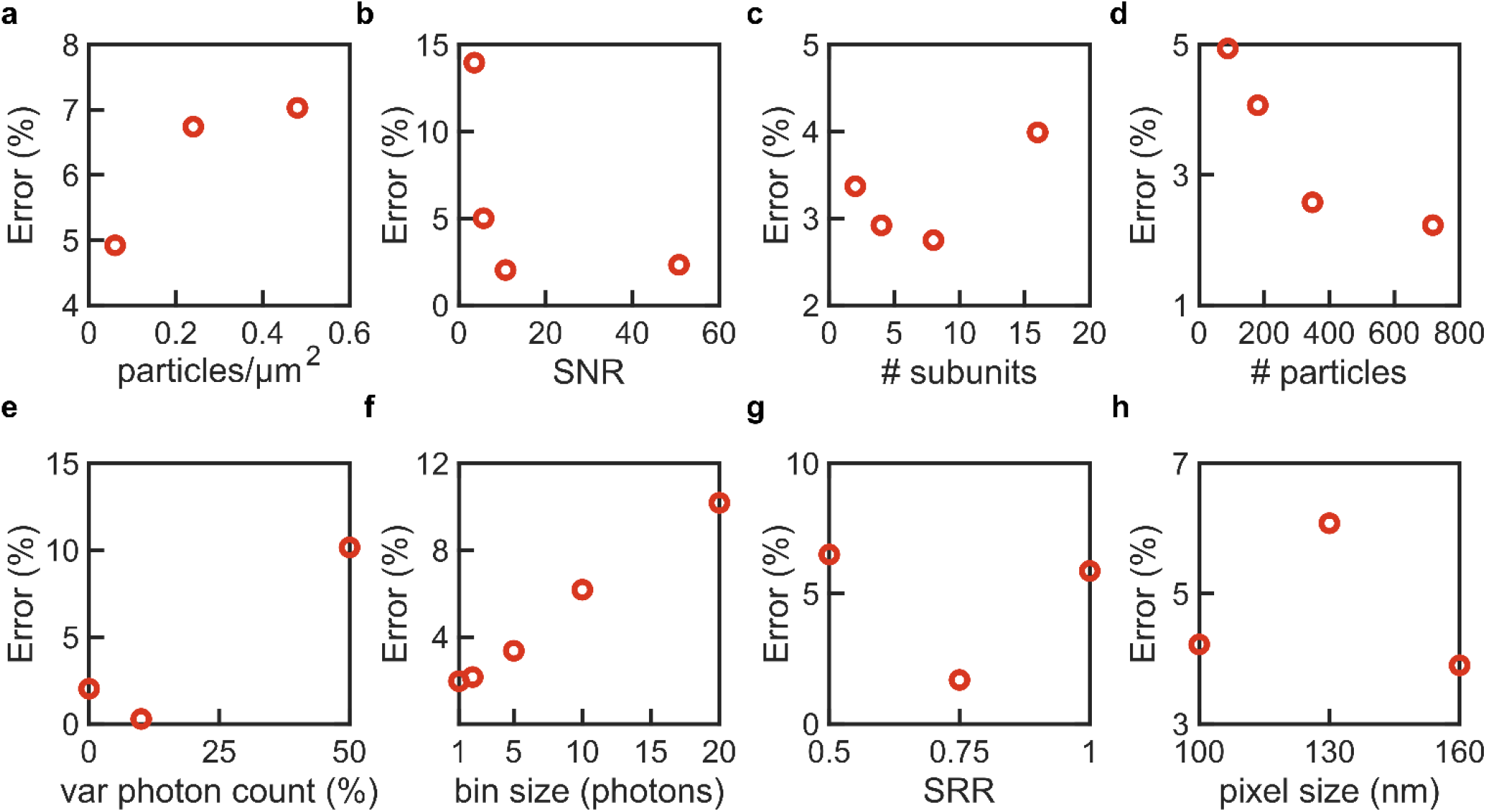
Assessment of the accuracy of subunit counting under different simulated experimental conditions. Measurement of the error against the **(a)** density of particles, **(b)** signal-to-noise ratio (SNR), **(c)** maximum number of subunits per complex, **(d)** number of monomeric particles selected for calibration, **(e)** inter complex variation in photon count, **(f)** kernel probability distribution function (pdf) bin size, **(g)** ratio of the sigma, or standard deviation, of the point spread function (PSF) of the underlying particles to ROI radius (SRR) and **(h)** pixel size. Base parameters used across all simulations (except for those varied): number of time frames = 500 frames, number of movies for calibration and unknown species (each) = at least 10 movies, maximum photon count = 10 photons, inter complex variation in photon count = 0%, sigma of the PSF of each complex = 130 nm, stoichiometry of the calibration species: (50% monomers, 25% dimers and 25% trimers) and stoichiometry of the unknown species: (50% monomers and 50% dimers). See **supplementary data** for camera parameters.

Our analysis shows that the bin size to generate a kernel pdf, as well as the number of calibration particles, may be a critical parameter to consider for fitting the intensity distributions, as increasing the bin size further increases the error rate whilst decreasing the number of calibration particles decreases the error rate. SAS employs a bin size of 5 photons, which provides an error of 3%, to ensure that the intensity information is not lost and the fitting procedure is not over-sensitive to fine fluctuations in the intensity curve. Expectedly, lowering the SNR to 3.54, which is remarkably low for single molecule experiments, affects the fidelity of the stoichiometry measurements, resulting in an error of 13.96%. A marginal improvement of the SNR to 5.68 yields a large improvement to the error (= 5.01%). Any improvements to the SNR beyond 5 to 10, yields diminishing returns on the error (> 2%). This result indicates that whilst the use of bright fluorophores and efficient detection setups are necessary to improve detection efficiency, beyond a certain point, is not necessarily correlated with improved counting accuracy by SAS. In contrast, our simulations indicate that intensity variation is a critical parameter to the accuracy of counting. Intensity variations greater than 25% can yield error values above 5%. These results favour the use of stable fluorophores that exhibit narrow emission spectrum and minimal photo blinking, as well as flat-field illumination schemes that minimize spatial variations in the excitation profile and unpolarized light as excitation source to ensure fluorophores under different orientations are equally excited.

Finally, the SRR affects the accuracy of counting. Surprisingly, however, our simulations indicate an optimal ratio of 0.75 at which the error is minimized to 1.69%. One possible explanation for this important finding is that for ROIs smaller than the full width of the particles the extracted intensities are inaccurate given that a large portion of the Point Spread Function (PSF) lies outside of the borders of the ROIs, yet, for ROIs much larger than the full width of the particles noise affects the extracted intensities. Our simulations point towards the importance of accurately measuring the mean standard deviation of the underlying particles in choosing the ROI radius.

### 2.3 Accuracy of counting with single molecule brightness analysis by SAS

Following, we assessed the accuracy of subunit counting for different stoichiometric configurations (**Figure 3a-i**). To do this, we simulated 9 different stoichiometric configurations with a maximum of 12 subunits where the proportion of species is constant (**Figure 3a-e**), decreasing (**Figure 3f** and **h**) or increasing (**Figure 3g** and **i**) with the number of subunits. The underlying species were also allowed to take monomeric up to hexameric units. Under all simulated configurations, the error did not exceed 15%. The accuracy of counting was particularly minimized when the proportion of species was held constant under all stoichiometries (**Figure 3a-e**). Although the error of the measurements was excellent throughout, we have noticed that, particularly where we have simulated increasing or decreasing proportions, a large fraction of the species was not recognized. This finding suggested that in a typical experiment, where not all single molecules assemble into higher order oligomers and where these oligomers add dimeric or higher-order units, larger complexes might not be recognized in the analysis.

**Figure 3.**
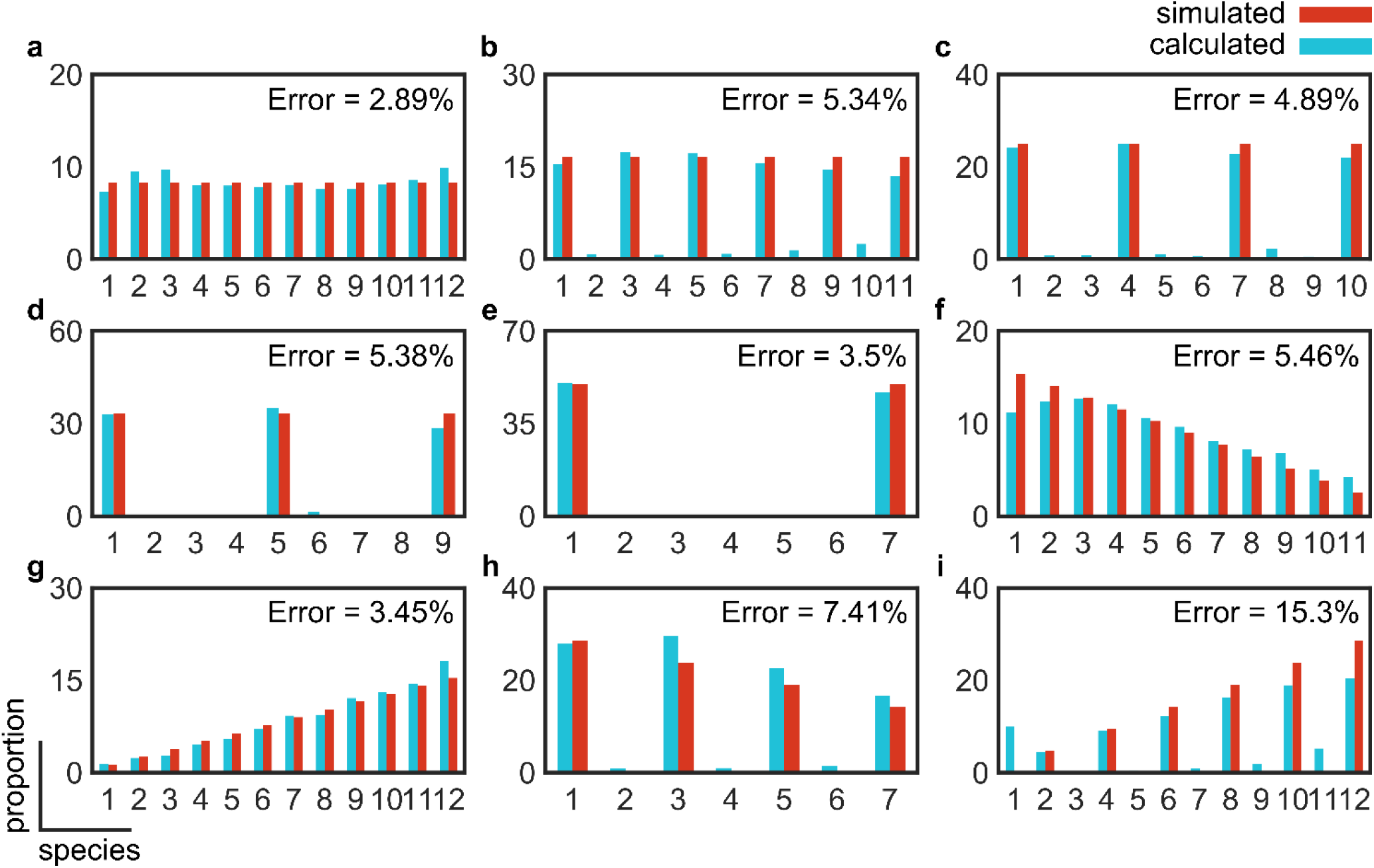
Assessment of the accuracy of subunit counting under different simulated stoichiometric configurations. Starting from a monomer as basic unit, we assessed the measurement of the error for equal proportions (8.33%) of **(a)** monomeric species, **(b)** dimeric species, **(c)** trimeric species, **(d)** tetrameric species, **(e)** hexameric species, **(f)** decreasing proportions (15.4% monomers, 14.1% dimers, 12.8% trimers, 11.5% tetramers, 10.3% pentamers, 9.0% hexamers, 7.7% heptamers, 6.4% octamers, 5.1% 9-mers, 3.8% 10-mers, 2.6% 11-mers and 1.3% 12-mers) based on monomeric units, **(g)** increasing proportions (same as **(f)** in reverse order) based on monomeric units, **(h)** decreasing proportions (28.6% monomers, 23.9% trimers, 19.0% pentamers, 14.3% heptamers, 9.5% 9-mers and 4.8% 11-mers) based on the addition of dimeric units and **(i)** increasing proportions (same as **(h)** in reverse order) based on the addition of dimeric units.

To investigate the reason behind the reduced performance of the software in quantifying the number of subunits in the mentioned stoichiometric configurations, we paid closer attention to the fittings of the idealized Gaussian mixture to the kernel density function of the unknown species. We found that marginal shifts in the mean intensity values of the calibration curves would propagate to high order oligomers beyond 8-to-10 subunits causing obvious misfits to the idealized mixture of Gaussians as is eluded to in **Figure 1g**. Furthermore, this issue could be more severe in a real, experimental setting where the intensity distribution of the underlying species might not follow the idealized Gaussian mixture due to imaging artefacts or photo quenching.

To solve this issue, we implemented a fitting refinement step where the mean intensity value of the calibration curve is scanned in a +/- 10 photons’ region, with 1 photon resolution, and the residual error is calculated after fitting with the Gaussian mixture model. The refined mean intensity value of the calibration curve is chosen where the residual error is minimum. Given that the mean intensity value of the monomer species is changed, we expect that the error would increase (i.e. be worsened) at the expense of recovering a larger number of species. Following this improvement, we first assessed two challenging configurations (**Figures 3f** and **h**). As expected, our assessments reveal an increase in the error from 5.46% to 7.99%, for the configuration based on the addition of monomers, and from 7.41% to 9.19% for the configuration based on the addition of dimers (**Figure 4a** and **b**). The advantages of using a refinement step were particularly observed in this last configuration where the number of recognized species increased from 7 to 11 out of a simulated 12 (**Figure 4b**). In all of the above, the SNR was set to 10.74 and the inter complex variation in photon count was set to 0% to ensure that none of these important photophysical parameters would complicate and affect our assessment of the accuracy of counting.

**Figure 4.**
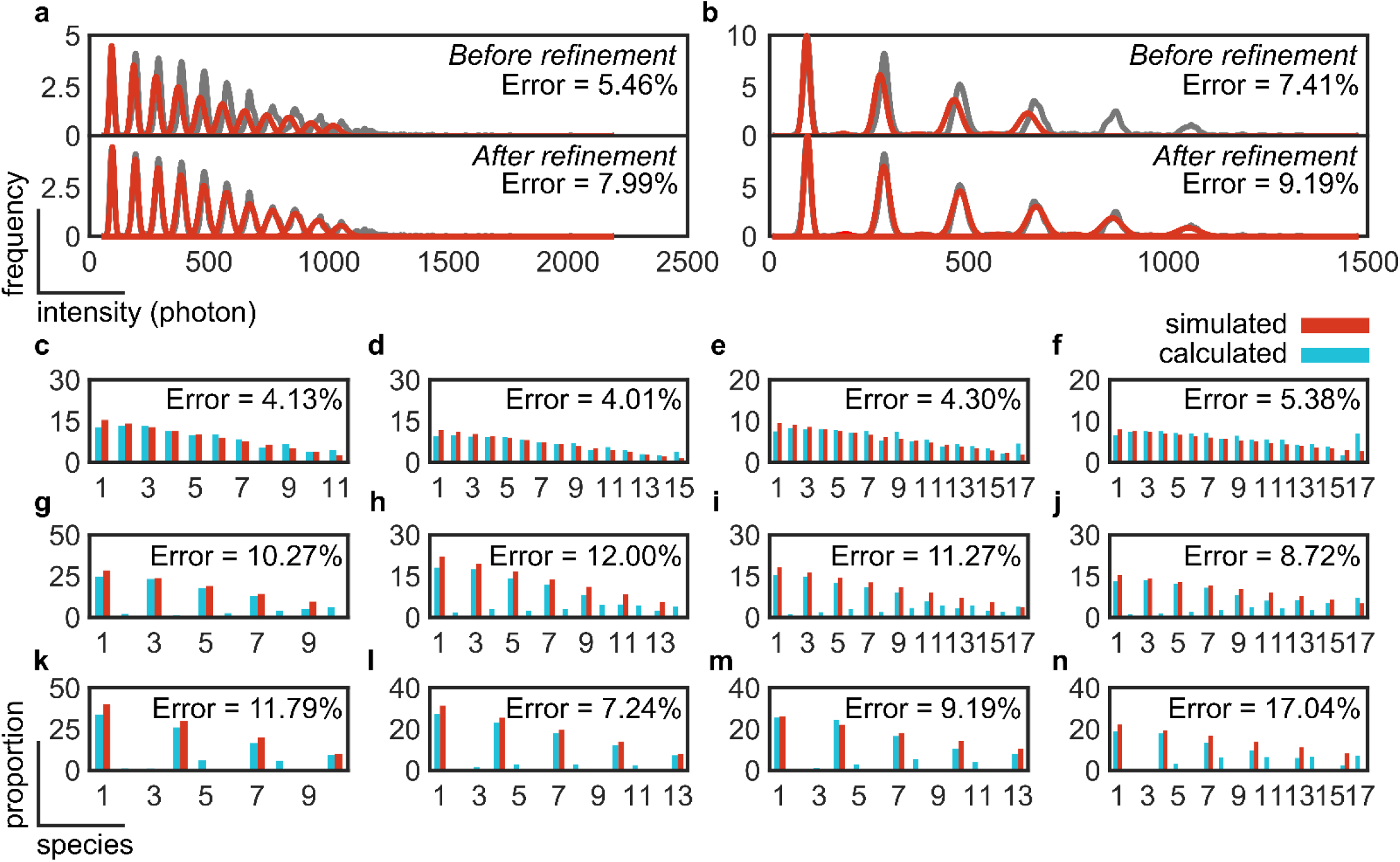
Assessment of the accuracy of subunit counting under real experimental conditions and challenging stoichiometric configurations. Kernel density distribution functions and error measurements for 12-mer stoichiometries based on the addition of **(a)** monomeric or **(b)** dimeric units before and after refinement simulated at an SNR of 10.74 (photon count = 10) and inter complex variation in photon count of 0%. Kernel density distributions are shown in grey and Gaussian mixture is shown in red. Error measurements based on the addition of **(c-f)** monomeric, **(g-j)** dimeric or **(k-n)** trimeric units of high order oligomers: **(c, g** and **k)**12-mer, **(d, h** and **l)** 16-mer, **(e, i** and **m)** 20-mer and **(f, j** and **n)** 24-mer. Simulations were done at an SNR of 5.68 (photon count = 5) and inter complex variation in photon count of 20%. Proportions of each species can be found in the **supplementary data**.

Next, we conducted a final round of assessment to establish the absolute limits of accurate counting with single molecule brightness analysis using the introduced refinement step for more challenging experimental conditions (i.e. an SNR of 5.68 and inter complex variation in photon count of 20%) and stoichiometric configurations (from 12- to 24-mers with decreasing proportion of species) (**Figures 4c-n**). On average, the error was lowest for the configurations based on monomeric, followed by dimeric and lastly trimeric units. All error measurements were below 15% except for the 24-mer in trimeric configuration where the measured error was 17.04%. Finally, and owing to the refinement step, high-order species were recognized in all cases, however, low proportions of species which were not simulated in the configurations based on the addition of dimeric and trimeric units were also produced.

Importantly, the end user needs to be aware that the refinement step helps to increase the accuracy at the expense of sensitivity. This step can be included or not in the analysis, as illustrated in the GUI (Fig. S1), thus leaving to the end-user the choice between accuracy or sensitivity, according to the specific experimental and analytical need.

In summary, this extensive analysis has shown the following:

1. The SNR and inter complex variation in photon count play an important role in dictating the accuracy as well as the number of recognised species within a complex of interest. Whilst marginal improvements to the SNR yield noticeable improvements to the accuracy of counting quickly followed by diminishing returns, inter complex variation in photon count has to be minimized at all times to maximize the accuracy of counting and number of recognized species.
2. The ROI size has to be optimized manually by the user in case the mean standard deviation in the PSF of the imaged complexes is known.
3. The stoichiometric occurrence does not affect the accuracy of measurements, but relevant factors are the basic unit (i.e. monomeric, dimeric or trimeric), and whether the proportion of species increases or decreases with the number of subunits.
4. Refining the mean intensity value of the calibration species can recover high-order species, but at the expense of a reduced counting accuracy as well as uncovering additional species which are absent in reality. The use of the refinement step is dependent on whether the user is interested in accurate or more comprehensive measurements of stoichiometry.

### 2.4 SAS evaluation on experimental biological data

Having extensively assessed the accuracy of counting with single molecule brightness analysis, we, finally, validated SAS on biological samples whose stoichiometry is known *a priori*, either as reported from structural studies or from prior subunit counting measurements performed manually. In addition, the chosen samples had to satisfy the following requirements:

1. The labelling efficiency had to be previously reported to ensure that any unlabelled species are accurately accounted for. Furthermore, the labelling efficiency had to be reported under exactly the same labelling conditions and using the same fluorophore as label in our experimental validation.
2. Highly compacted structures, in which the underlying fluorophores are closely located from one another, were avoided. In doing so, we wanted to ameliorate the hard to simulate effects of fluorophore quenching on the measured intensity of the complex of interest.
3. For validation purposes, the complex of interest would be known to take stable stoichiometries which would not change under the course of an experiment or under slightly different conditions. This is to ensure that our results would, to the best of our knowledge, match those reported.
4. The complex of interest has to be assembled from its individual components *in vitro* and *in situ*. Oligomeric complexes which can only be imaged inside their physiological, cellular environment were excluded as their densities, as well as the behaviour of the host cellular system, cannot be appropriately controlled.

We identified the Bcl-2-associated-X-protein (BAX), which is known to assemble into multiple species based on dimer units^17^, and the lipid scramblase Atg9, which has been recently reported to assemble as a homotrimer^28-30^, as candidate systems. To this end, we reconstituted labelled BAX oligomers into a Supported Lipid Bilayer (SLB) and imaged them under a TIRF microscope (see **methods**). We then compared the proportion of species measured using SAS with those measured manually as reported in^17^ (**Figure 5a-e** and **Supplementary Figure 3**). Our measurements show excellent agreement with that reported, revealing the dimeric stoichiometry of BAX. Moreover, owing to the entirely automated pipeline of SAS, it took us 3 minutes to process one dataset which typically took hours to days, to process through the manual selection of clean traces, as well as optimization of detection under various experimental conditions. Similarly, stoichiometry experiments on Atg9 complexes processed by SAS showed excellent agreement with the literature data, with Atg9 assembling predominantly as a trimer (with minor high order aggregates/complexes based on trimer units, **Figure 5f**).

**Figure 5.**
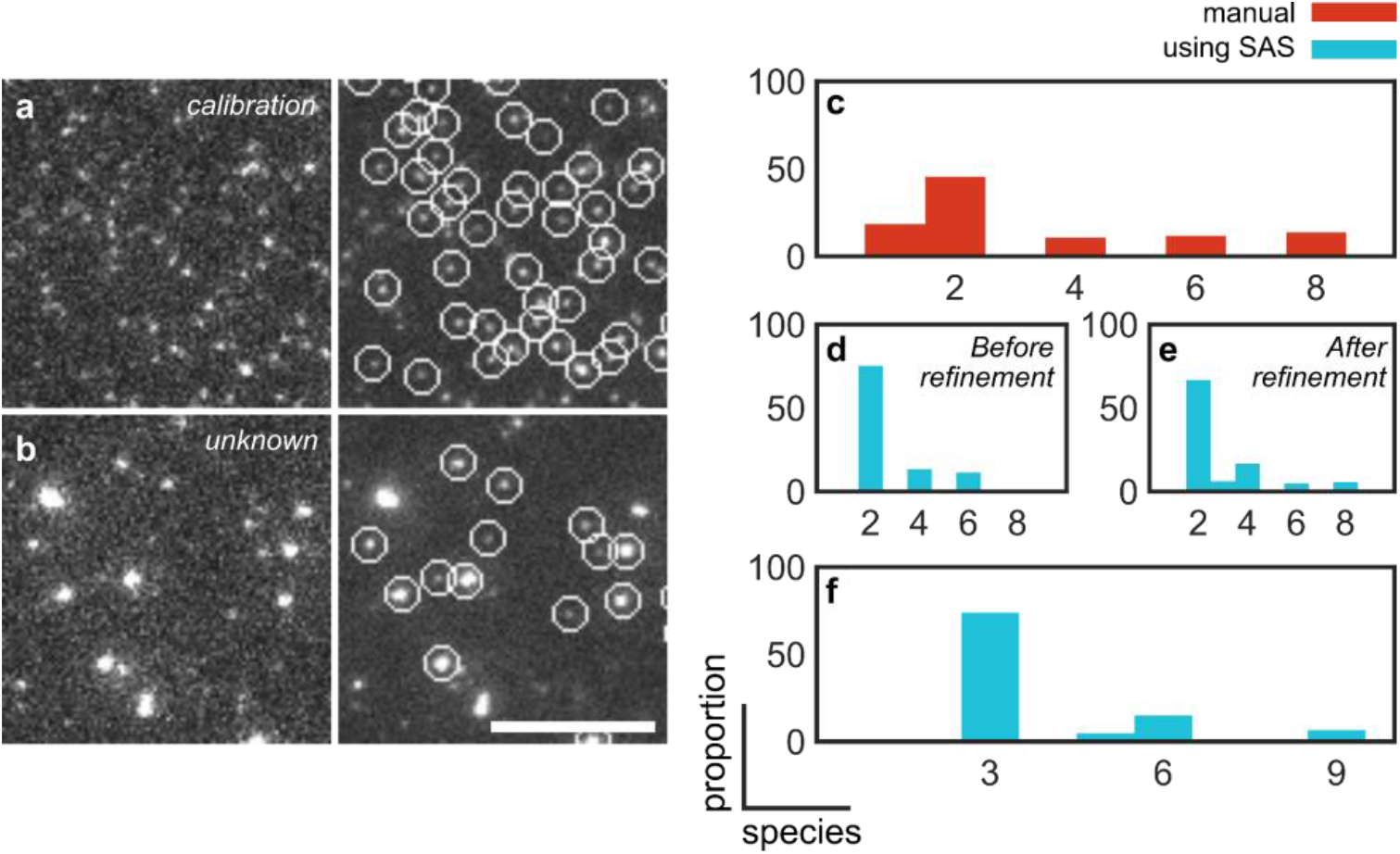
Comparison of subunit counting accuracy with a manual (semi-automated) pipeline and SAS applied on experimental data. **(a)** A calibration dataset of monomeric Bax labelled with ATTO 488 dye. **(b)** A set of unknown stoichiometries of labelled BAX molecules. Scale bar, 5 µm. Subunit counting of BAX performed **(c)** manually and using SAS **(d)** before and **(e)** after refinement. **(f)** Subunit counting of Atg9 performed using SAS.

## CONCLUSIONS

In summary, here we have systematically and extensively assessed the accuracy of subunit counting using brightness analysis. We have established the experimental conditions, as well as assessed complex stoichiometric configurations, under which this method can count with accuracies exceeding 85%. Our analysis serves as an important resource for experimentalists in need of accurately counting the copy number of proteins in a variety of stoichiometric configurations and under a wide range of challenging experimental conditions. In order to perform this analysis, we developed a fully automated computational pipeline that is simple to use and which serves as a fundamental tool for future experiments of this type. We expect our analysis, and software, to empower the use of optical microscopy in structural studies of complex, large and heterogenous macromolecular assemblies with single molecule sensitivity.

## Methods

### Simulating ground-truth data

To simulate the ground-truth data, a pre-set number of particles (i.e. complexes) were randomly scattered across images which are 256 by 256 pixels in size (except for those simulated in **Figures 4c-n** which were 1024 by 2024 pixels in size to accommodate for the larger number of complexes in the same field of view without significantly increasing the density) with pixel size of 100 nm. The standard deviation in the PSF of each particle was fixed at 130 nm. Each complex contained a pre-set number of molecules with intensities sampled from a random number generator based on a normal distribution with a mean intensity of *I*_*m*_ and standard deviation of *v*, where *v* is the inter complex variation in the intensity. Each molecule had a photobleaching time that was randomly sampled from 1 to the maximum number of frames which in our simulation was set to 500 frames. The intensity of each molecule was set to the sampled intensity before the photobleaching time point and to 0 afterwards. Each molecule was convoluted with a 2D Gaussian Kernel. To account for the noise statistics of Electron-Multiplying Charge Coupled Detectors (EMCCDs), the produced images in counts *I*_*c*_ were modified as follows:

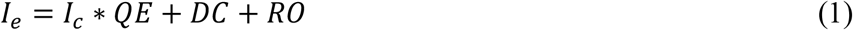

where *I*_*e*_ is the image signal in electrons, *QE* is the quantum efficiency, *DC* is the dark current and *RO* is the readout noise. The quantum efficiency was set at 95%, dark current was set at 0.0002 electrons/second and readout noise was set to 1 electron. These values are typical of commercially available EMCCDs. To simulate noise, *I*_*e*_ was modified as follows:

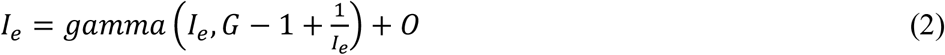

where *G* is the camera gain (corrected for conversion factor) and *O* is the camera bias offset. The gain was set to 58.8 (electron gain of 300 and conversion factor of 5.1) and offset to 100. This noise model was chosen to best replicate the electron-multiplication feature in EMCCDs^26^. The produced images were saved in the big TIFF format at 16 bits for further processing using SAS.

### Detecting single molecule using a deep convolutional network

Detection of each complex / molecule was carried out as described in^25^. Briefly, we developed and used DeepSinse, a simple, multi-layer Convolutional Neural Network (CNN) architecture to enable fast detection of single molecules using as few parameters as possible. Our neural network is composed of a CNN, a dense layer and a SoftMax (classification) layer. The neural network was first trained to classify simulated ground-truth datasets of noise and Gaussian bursts in pre-labelled Regions Of Interest (ROIs), then validated on different, unseen, datasets of pre-labelled ROIs. We then tested it on ground-truth generated ROIs. The neural network is finally deployed by feeding an image into a peak-finding algorithm based on identifying regional maxima. The peak-finding algorithm outputs hundreds of noise- and burst- containing ROIs which are then fed into the trained network for classification, thus, resulting in an annotated image. Burst-containing, and pure-noise, images were simulated with the formers’ peak burst intensities varying from 50 to 100 counts corresponding to signal-to-noise ratios from 24.85 to 45.81 and 100 images were simulated with pure noise. Pre-annotated ROIs were picked from each of these images, intensity scaled between 0 and 1 to avoid subjective segmentation parameters such as the intensity threshold, shuffled and fed into the neural network for training. To optimize performance, the neural network was trained using different ROI radii and number of ROIs. The lowest false negative rate FNR (61.45%) and false positive rate FPR (0.3%) were achieved at a ROI radius of 5 pixels and 10,000 training ROIs. The trained network was then integrated into SAS for immediate use. Particles that are less than 6 pixels away from the borders of each image, as well as those which are less than 5 pixels apart from each other are rejected. Finally, the detected particles were fed into a least-squares solver to fit a Gaussian function to the PSF of each particle using the cpufit plugin^27^ and extract a value to the standard deviation which was compared to a user-specified value (200 nm for the simulated and experimental data as commonly used in single molecule microscopy^5^). Particles exhibiting a standard deviation value smaller than the user-set value are accepted and, otherwise, rejected. This approach allows for discarding multiple particles present in the same ROI.

### Extraction of intensity traces

The intensity of the detected and accepted molecules were extracted from the movies by drawing ROI with a user-specified radius around each of these molecules at each time point. The local background at each time point is calculated by extracting the intensity in a region around the ROI, 2 pixels larger than the ROI. Intensity traces are used for both, the selection of monomeric complexes for calibration (see “Selection of monomeric traces for calibration”), and for the extraction of the intensity value for the detected molecule. In this last case, the maximum intensity of each trace is calculated by taking the median of the first 5 time points of the intensity trace, and the background is subtracted by taking the median of the intensity of the last 5 time points of the background trace (which, theoretically, should yield the same result as the first 5 time points).

### Selection of monomeric traces for calibration

The traces extracted for particles belonging to the calibration sample are fed into a trace annotator which calculates the absolute gradient of each intensity trace, normalizes the calculated gradient between 0 and 1 and, finally, extracts the number of peaks in the normalized gradient above a threshold value, known as the minimum peak height, which we set at 0.5 (for simulated data), 0.9 (for experimental data). The value of the minimum peak height was optimized once for each dataset by inspection of the calibration curve so that, ideally, a single peak corresponding to the monomeric traces was dominant. Gradients with a single peak were chosen as monomeric traces (i.e. arising from a complex with a single molecule).

### Calculating the proportion of labelled species

After measuring the intensity for each detected and accepted particle (measured as described in “Extraction of intensity traces”), a kernel probability distribution function (pdf) with a bandwidth of 5 photons was calculated from the intensity values of the calibration species and normalized between 0 and 1. A peak finder was subsequently used to find peaks in the normalized kernel pdf with a minimum height of 0.8 (normalized value). The first peak, corresponding to the intensity distribution of monomers, was selected and its mean intensity value (*I*_*m*_) was used to fit the following Gaussian Mixture Model (GMM) using a non-linear curve fitting solver to the kernel pdf to extract the standard deviation in the intensity values of the population of monomers (*σ*_*m*_):

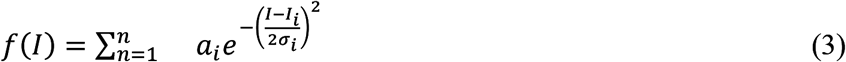

Where *a*_*i*_ *I*_*i*_ *and σ*_*i*_ the amplitude, mean intensity value and standard deviation in intensity value of each Gaussian. *n* is the number of Gaussians and was set to 2 to account for any dimers present in the sample used for calibration. In this case, 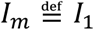 and 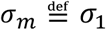. Following, a kernel pdf was constructed from the sample of unknown species with a bandwidth of 5 photons which was fit, using a non-linear curve fitting solver, with an idealized GMM:

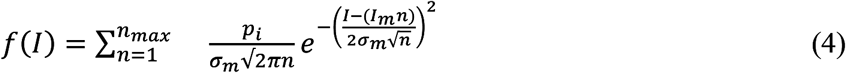

Where *p*_*i*_ is the proportion of each species corresponding to the area below each Gaussian curve and *n*_*max*_ is the maximum number of Gaussians. *n*_*max*_ was calculated as described in^22^ from *I*_*m*_ and *σ*_*m*_ using the following formula:

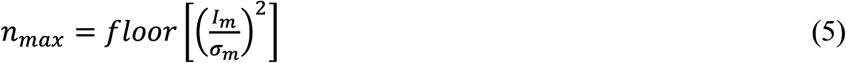

Constraining the maximum number of Gaussians as such does not prevent overfitting. To account for overfitting, we performed non-linear curve fitting with *n* from 1 to *n*_*max*_. In each time we performed a fit, we calculated the root sum squared residual and whenever this was less than 95% of the minimum calculated value, this last was set as the minimum value and the used number of Gaussians as *n*_*max*_. Finally, an optional refinement step was performed were *I*_*m*_ was scanned in a ± 10 photons region with 1 photon resolution and modified to where the root sum squared residual was minimized.

The residual error is calculated in an ascending order of Gaussians. The minimum residual error is also calculated in that specific order. For configurations where the proportion of species decreases with increasing the species number (as in figure 3h) a minimum residual error is reached with low species numbers which have higher proportions. This is not the case with configurations where the proportion of species increases with increasing the specie number (as in figure 3i).

### Correcting for labelling efficiency to calculate the true proportion of labelled and unlabelled species

Labelling efficiency was corrected for as described in^17^. Briefly, each molecule was assumed to be either labelled (1) or not labelled (0). To uncover the true proportion of species from those measured above, we constructed a binomial probability distribution function of the following form:

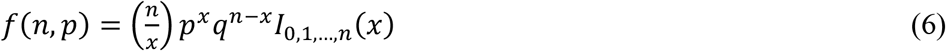

Where *x* is the species number, *n* is the number of trials, *p* is the probability of a molecule being labelled (set to the labelling efficiency) and *q* is the probability of a molecule not being labelled (= 1 − *p*). *x* and *n* took values from 1 to the number of measured species. A linear least-squares problem solver was then used to calculate the true proportion of species taking the constructed binomial probability distribution function as the multiplier matrix and the measured proportion of species as the constant vector. The solver was constrained between 0 and 100% across all species.

### Preparing BAX reference sample and imaging setup used for experimental validation

The sample was prepared as described in^17^. Briefly, egg phosphatidylcholine and cardiolipin (Avanti Polar Lipids, US) were mixed in a 7:3 ratio and dissolved in chloroform that was evaporated under reduced pressure for 3 hours. The lipid film was then resuspended with 150 mM NaCl, 10 mM Hepes (pH7.4) to a final concentration of 1 mg/mL. The lipid solution was subjected to five cycles of freezing and thawing after which they were manually extruded through a polycarbonate membrane with a defined pore size (100 nm) using glass syringes. The formed large unilamellar vesicles (LUVs) were then incubated with 10 nM Atto488 labelled Bax (labelling efficiency 84%) for 1 hour at 43°C to give proteoliposomes which were subsequently diluted 1:10 with untreated liposomes. The supported lipid bilayer (SLB) was formed by incubating the diluted proteoliposomes on a glass slide, previously cleaned with a piranha solution, at 37 °C for 2 minutes in the presence of 3 mM CaCl_2_ and then washed several times with 150 mM NaCl, 10 mM Hepes (pH7.4) to remove non-fused vesicles. Samples were imaged on the setup described below for a total of 1200 frames under a 35 ms exposure time and 25 ms delay between frames with a power density of ∼ 1 kW/cm^2^.

The sample was imaged on a Total Internal Reflection Fluorescence (TIRF) microscope. The laser excitation from a 4-wavelengths (405 nm, 488 nm, 561 nm and 647 nm) laser engine (iChrome MLE, Toptica Photonics AG, DE) was coupled into a multi-mode fibre onto a TIRF-alignment module (Laser TIRF 3, Carl Zeiss AG, DE) inserted into the side port of an upright microscope (Axiovert 200, Carl Zeiss AG). Using the TIRF- alignment module, the excitation light was focused onto the back focal plane of a 100x, 1.46 Numerical Aperture (NA) objective (Apo-TIRF, Carl Zeiss AG) after passing through a triple-band clean-up filter (TBP 483 + 564 + 642 (HE), Carl Zeiss AG) and being reflected off a quad-band dichroic filter (TFT 506 + 582 + 659 (HE), Carl Zeiss AG). Image was additionally magnified by 1.6x to obtain a final pixel size of 100 nm. The emission was collected using the same objective, passed through a triple-band emission filter (TBP 526 + 601 + 688 (HE), Carl Zeiss AG) and focused on an EMCCD camera (iXon 88X, Andor, IE) cooled at -70 degC.

### Preparing Atg9 reference sample and imaging setup used for experimental validation

Atg9 was overexpressed in yeast as GFP-3xFLAG fusion protein and purified as previously described^31^. For the generation of Atg9 containing liposomes or protein free liposomes, 50% DOPC, 30% DOPE and 20% DOPS (Avanti Polaris) were mixed and dried under vacuum 1h at 37°C. The lipid film was resuspended in buffer A (300 mM KCl, 50 mM HEPES, pH 7.4) to a concentration of 20 mg/ml and sonicated 15 min. Generated SUVs were destabilized by adding 35 mM CHAPS followed by incubation 1h at 23°C. 1.5mg of lipids were mixed with 10 µg of the purified protein or with lysis buffer (for protein free liposomes) and the mixes were incubated 1h at 4°C. Then, samples were diluted 10 times in buffer A to reduce the concentration of detergent below the critical micelle concentration (CMC). Samples were dialyzed using Slide-A-Lyzer dialysis cassettes 20 MWCO (Thermo Scientific) against buffer A plus 0.2 g of Bio-beads SM2 adsorbent Media (BIO-RAD) per liter of buffer. Reconstituted liposomes were freeze-thawed two times. The SLB was formed by incubating 1:10 diluted proteoliposomes on a glass slide, cleaned with a piranha solution, at 37°C for 10min with 3mM CaCl_2_, and washed 15 times with 150 mM NaCl, 10mM Hepes (pH7.4) buffer to remove non-fused vesicles.

Samples were imaged on a custom-designed TIRF microscope for a total of 2000 frames under a 30 ms exposure time. Laser excitation from a 488 nm laser, max. power 400 mW (Sapphire, Coherent) was coupled into a single mode polarization maintaining fiber to a TIRF module connected to an Olympus IX83 inverted microscope with hardware autofocus system (IX3-ZDC, Olympus) and 100x oil-immersion objective ((UPLAPO100xOHR). Image was additionally magnified by 1.6x (IX3-CAS, Olympus) to obtain a final magnification of 160x and a pixel size of 100 nm. Fluorescence was filtered by a four-line polychroic mirror (zt405/488/561/640rpc, Chroma, 3 mm) and rejection band filter (zet405/488/561/647 TIRF, Chroma), and the emission was focussed on an iXon Ultra EMCCD Camera (Andor Technologies).

## Data availability

The data that support the findings of this study are available as Supplementary Data.

## Code availability

Updated versions of the source code for SAS, as well as guiding instructions, can be obtained from https://github.com/jdanial/SAS. A compilation of SAS for Windows OS is available as Supplementary Software.

## Contributions

K.C. and A.J.G.S. conceived the study. J.S.H.D., Y.Q., A.J.G.S. and K.C designed the study with the contribution of U.R. and R.S. K.C. performed BAX experiments. E.G.M. and S.C. performed Atg9 experiments. J.S.H.D. and Y.Q. wrote the software. K.C. performed the manual analysis. J.S.H.D. and E.G.M. performed the software analysis. K.C, J.S.H.D., U.R. and R.S. assessed performance. K.C., C.U. and A.J.G.S. provided supervision and infrastructures. All authors contributed to writing the manuscript.

## Acknowledgments

This work was supported by a research associateship from King’s College, University of Cambridge awarded to J.S.H.D, a scholarship from the International Max-Planck Research School and the University of Tubingen awarded to R.S, a Deutsche Forschungsgemeinschaft (DFG) grant UN111/13-1 awarded to C.U., a DFG GA164/3-1, starting (grant no. 309966) and consolidator (APOSITE) ERC grants awarded to A.J.G.S., and an Elite Postdoctoral Fellowship from the Baden-Wuerttemberg Foundation and DFG grant INST 190/194-1 awarded to K.C.

