## Supplementary material for "Systematic assessment of the accuracy of subunit counting in biomolecular complexes using automated single molecule brightness analysis": Supplementary information.docx

**Supplementary Figure 1**

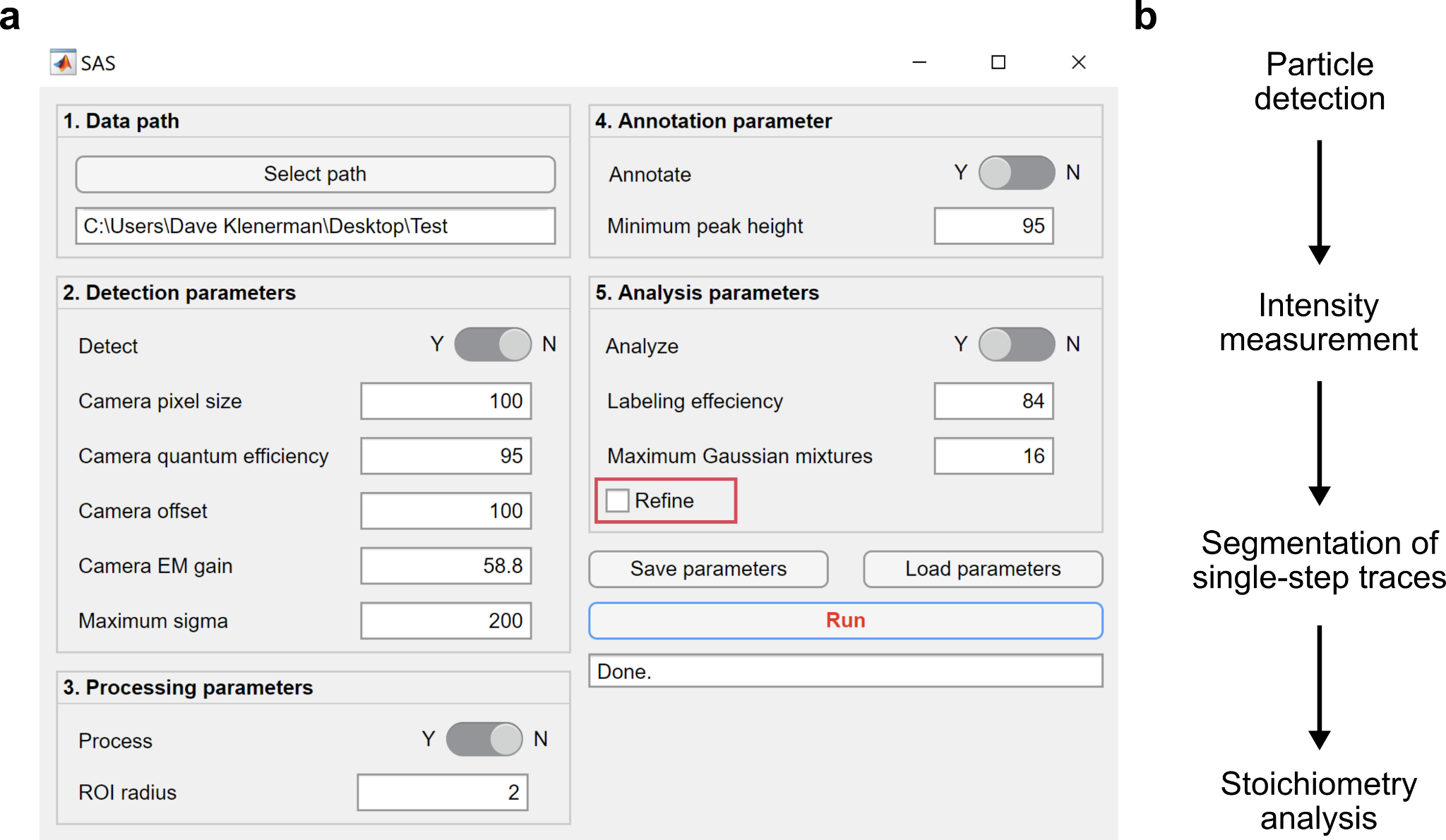

**Figure S1.** (a) Graphical user interface (GUI) of SAS. The GUI is divided into five panels: 1. Data import; 2. Detection parameters to be fed with the experimental parameters used by the experimenter during data acquisition; 3. Processing parameters that allows the user to select the ROI radius based on the experimental parameters used; 4. Annotation of parameters in a text file and choice of the threshold value (minimum peak height) for appropriate monomeric curves selection; 5. Analysis parameters that allow to introduce the values of the protein labelling efficiency and of the maximum number of Gaussian mixtures the user aims to resolve. Importantly, the refinement step can be selected in this panel (red box). (b) SAS workflow.

**Supplementary Figure 2**

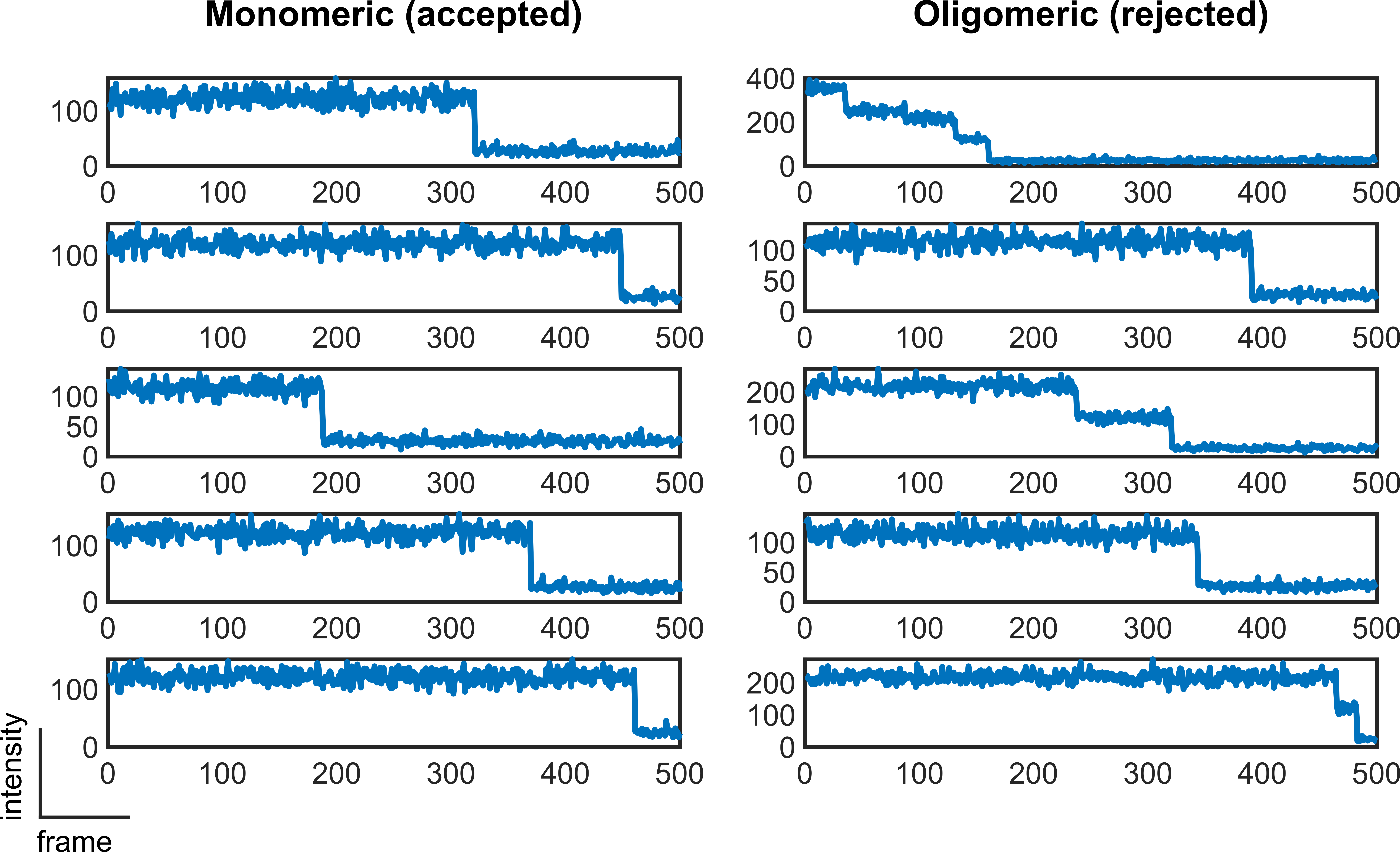

**Figure S2.** Gallery of randomly accepted (monomeric) and rejected (oligomeric) traces selected by the trace annotator tool of SAS.

**Supplementary Figure 3**

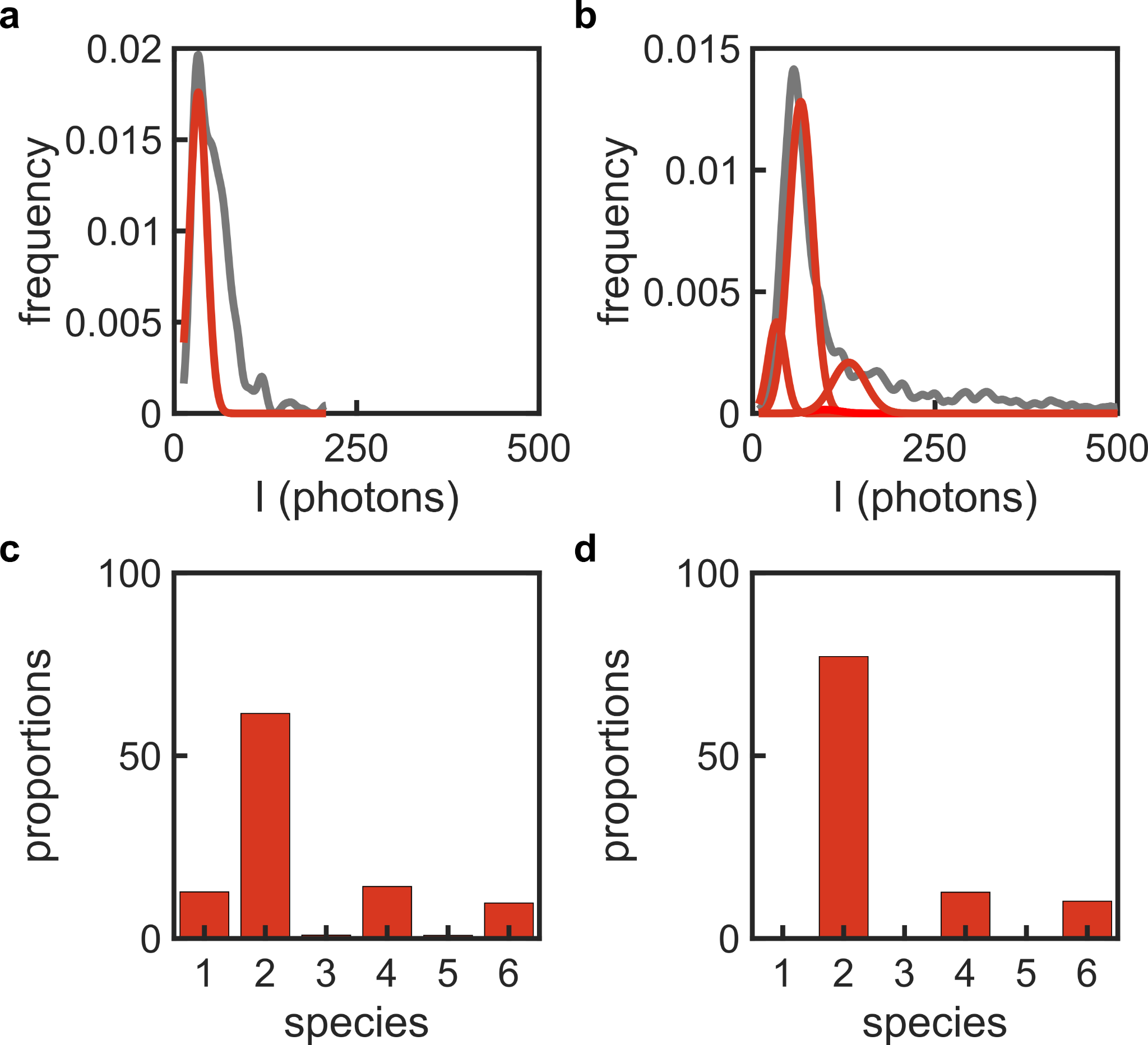

**Figure S3.** Graphs of the (a) calibration, and (b) unknown (c) intensity distributions obtained from the experimental data to measure the stoichiometry of BAX complexes. Percentage of occurrence of Bax species (c) before, and (d) after labelling correction (labelling efficiency 84%). Illustrated data are without refinement. Results after refinement are shown in figure 5.

**Supplementary Table 1**

| **Figure** | **Stoichiometric configuration**  **Experimental conditions (if applicable)**  **Reference in supplementary data / reference dataset** |
| --- | --- |
| Figure 2a | Calibration: 1-mer (50%), 2-mer (25%), 3-mer (25%) [varied from 20 to 320 particles per movie]  Unknown: 1-mer (50%), 2-mer (50%) [varied from 20 to 320 particles per movie] |
|  | # frames = 500, # movies = 10, max. photon count = 10, variation in photon count = 0%, lateral sigma = 130 nm, frame size = 256 pixels x 256 pixels |
|  | Columns A to O / dataset # 2 to 5 |
| Figure 2b | Calibration: 1-mer (50%), 2-mer (25%), 3-mer (25%) [80 particles per movie]  Unknown: 1-mer (50%), 2-mer (50%) [80 particles per movie] |
|  | # frames = 500, # movies = 10, max. photon count = 2 to 50, variation in photon count = 0%, lateral sigma = 130 nm, frame size = 256 pixels x 256 pixels |
|  | Columns P to AI / dataset # 6 to 10 |
| Figure 2c | Calibration: 1-mer (50%), 2-mer (25%), 3-mer (25%) [80 particles per movie]  Unknown: varied from 1-mer (50%), 2-mer (50%) [80 particles per movie] to 1-to-16-mers (3.125% each) |
|  | # frames = 500, # movies = 10, max. photon count = 10, variation in photon count = 0%, lateral sigma = 130 nm, frame size = 256 pixels x 256 pixels |
|  | Columns AJ to BC / dataset # 11 to 14 |
| Figure 2d | Calibration: 1-mer (50%), 2-mer (25%), 3-mer (25%) [80 particles per movie]  Unknown: 1-mer (50%), 2-mer (50%) [80 particles per movie] |
|  | # frames = 500, # movies = varied from 5 to 40, max. photon count = 10, variation in photon count = 0%, lateral sigma = 130 nm, frame size = 256 pixels x 256 pixels |
|  | Columns BD to BW / dataset # 15 to 18 |
| Figure 2e | Calibration: 1-mer (50%), 2-mer (25%), 3-mer (25%) [80 particles per movie]  Unknown: 1-mer (50%), 2-mer (50%) [80 particles per movie] |
|  | # frames = 500, # movies = 10, max. photon count = 10, variation in photon count = 0% to 50%, lateral sigma = 130 nm, frame size = 256 pixels x 256 pixels |
|  | Columns BX to CL / dataset # 19 to 22 |
| Figure 2f | Calibration: 1-mer (50%), 2-mer (25%), 3-mer (25%) [80 particles per movie]  Unknown: 1-mer (50%), 2-mer (50%) [80 particles per movie] |
|  | # frames = 500, # movies = 10, max. photon count = 10, variation in photon count = 0%, lateral sigma = 130 nm, frame size = 256 pixels x 256 pixels. Analysis bin sized varied from 1 to 20. |
|  | Columns HP to IR / dataset # 45 to 49 |
| Figure 2g | Calibration: 1-mer (50%), 2-mer (25%), 3-mer (25%) [80 particles per movie]  Unknown: 1-mer (50%), 2-mer (50%) [80 particles per movie] |
|  | # frames = 500, # movies = 10, max. photon count = 10, variation in photon count = 0%, lateral sigma = varied from 100 nm to 200 nm, frame size = 256 pixels x 256 pixels |
|  | Columns CM to DA / dataset # 23 to 25 |
| Figure 2h | Calibration: 1-mer (50%), 2-mer (25%), 3-mer (25%) [80 particles per movie]  Unknown: 1-mer (50%), 2-mer (50%) [80 particles per movie] |
|  | # frames = 500, # movies = 10, max. photon count = 10, variation in photon count = 0%, lateral sigma = 130 nm, frame size = 256 pixels x 256 pixels, pixel size varied from 100 nm to 160 nm. |
|  | Columns IT to JJ / dataset # 50 to 52 |
| Figure 3a | Calibration: 1-mer (50%), 2-mer (25%), 3-mer (25%) [80 particles per movie]  Unknown: 1-to-12-mers (8.33% each) [96 particles per movie] |
|  | # frames = 500, # movies = 10, max. photon count = 10, variation in photon count = 0%, lateral sigma = 130 nm, frame size = 256 pixels x 256 pixels |
|  | Columns DB to DF / dataset # 26 |
| Figure 3b | Calibration: 1-mer (50%), 2-mer (25%), 3-mer (25%) [80 particles per movie]  Unknown: 1,3,5,7,9 and 11-mers (16.67% each) [96 particles per movie] |
|  | # frames = 500, # movies = 10, max. photon count = 10, variation in photon count = 0%, lateral sigma = 130 nm, frame size = 256 pixels x 256 pixels |
|  | Columns DG to DK / dataset # 27 |
| Figure 3c | Calibration: 1-mer (50%), 2-mer (25%), 3-mer (25%) [80 particles per movie]  Unknown: 1,4,7 and 10-mers (25% each) [96 particles per movie] |
|  | # frames = 500, # movies = 10, max. photon count = 10, variation in photon count = 0%, lateral sigma = 130 nm, frame size = 256 pixels x 256 pixels |
|  | Columns DL to DP / dataset # 28 |
| Figure 3d | Calibration: 1-mer (50%), 2-mer (25%), 3-mer (25%) [80 particles per movie]  Unknown: 1,5 and 9-mers (33.33% each) [96 particles per movie] |
|  | # frames = 500, # movies = 10, max. photon count = 10, variation in photon count = 0%, lateral sigma = 130 nm, frame size = 256 pixels x 256 pixels |
|  | Columns DQ to DU / dataset # 29 |
| Figure 3e | Calibration: 1-mer (50%), 2-mer (25%), 3-mer (25%) [80 particles per movie]  Unknown: 1 and 7-mers (50% each) [96 particles per movie] |
|  | # frames = 500, # movies = 10, max. photon count = 10, variation in photon count = 0%, lateral sigma = 130 nm, frame size = 256 pixels x 256 pixels |
|  | Columns DV to DZ / dataset # 30 |
| Figure 3f | Calibration: 1-mer (50%), 2-mer (25%), 3-mer (25%) [80 particles per movie]  Unknown: 1-mer (15.38%), 2-mer (14.10%), 3-mer (12.82%), 4-mer (11.54%), 5-mer (10.26%), 6-mer (8.97%), 7-mer (7.69%), 8-mer (6.41%), 9-mer (5.13%), 10-mer (3.85%), 11-mers (2.56% each) and 12-mer (1.28%) [312 particles per movie] |
|  | # frames = 500, # movies = 10, max. photon count = 10, variation in photon count = 0%, lateral sigma = 130 nm, frame size = 256 pixels x 256 pixels |
|  | Columns EA to EE / dataset # 31 |
| Figure 3g | Calibration: 1-mer (50%), 2-mer (25%), 3-mer (25%) [80 particles per movie]  Unknown: 1-mer (1.28%), 2-mer (2.56%), 3-mer (3.85%), 4-mer (5.13%), 5-mer (6.41%), 6-mer (7.69%), 7-mer (8.97%), 8-mer (10.26%), 9-mer (11.54%), 10-mer (12.82%), 11-mers (14.10% each) and 12-mer (15.38%) [312 particles per movie] |
|  | # frames = 500, # movies = 10, max. photon count = 10, variation in photon count = 0%, lateral sigma = 130 nm, frame size = 256 pixels x 256 pixels |
|  | Columns EF to EJ / dataset # 32 |
| Figure 3h | Calibration: 1-mer (50%), 2-mer (25%), 3-mer (25%) [80 particles per movie]  Unknown: 1-mer (28.57%), 3-mer (23.81%), 5-mer (19.10%), 7-mer (14.29%), 9-mer (9.52%) and 11-mer (4.76%) [168 particles per movie] |
|  | # frames = 500, # movies = 10, max. photon count = 10, variation in photon count = 0%, lateral sigma = 130 nm, frame size = 256 pixels x 256 pixels |
|  | Columns EK to EO / dataset # 33 |
| Figure 3i | Calibration: 1-mer (50%), 2-mer (25%), 3-mer (25%) [80 particles per movie]  Unknown: 2-mer (4.76%), 4-mer (9.52%), 6-mer (14.29%), 8-mer (19.05%), 10-mer (23.81%) and 12-mer (28.57%) [168 particles per movie] |
|  | # frames = 500, # movies = 10, max. photon count = 10, variation in photon count = 0%, lateral sigma = 130 nm, frame size = 256 pixels x 256 pixels |
|  | Columns EP to ET / dataset # 34 |
| Figure 4a | Calibration: 1-mer (50%), 2-mer (25%), 3-mer (25%) [80 particles per movie]  Unknown: 1-mer (15.38%), 2-mer (14.10%), 3-mer (12.82%), 4-mer (11.54%), 5-mer (10.26%), 6-mer (8.97%), 7-mer (7.69%), 8-mer (6.41%), 9-mer (5.13%), 10-mer (3.85%), 11-mers (2.56% each) and 12-mer (1.28%) [312 particles per movie] |
|  | # frames = 500, # movies = 10, max. photon count = 5, variation in photon count = 20%, lateral sigma = 130 nm, frame size = 1,024 pixels x 1,024 pixels |
|  | Columns EV to EZ (rows 1 to 14) / dataset # 31 copy |
| Figure 4b | Calibration: 1-mer (50%), 2-mer (25%), 3-mer (25%) [80 particles per movie]  Unknown: 1-mer (28.57%), 3-mer (23.81%), 5-mer (19.10%), 7-mer (14.29%), 9-mer (9.52%) and 11-mer (4.76%) [168 particles per movie] |
|  | # frames = 500, # movies = 10, max. photon count = 5, variation in photon count = 20%, lateral sigma = 130 nm, frame size = 1,024 pixels x 1,024 pixels |
|  | Columns FB to FF (rows 1 to 14) / dataset # 33 copy |
| Figure 4c | Calibration: 1-mer (50%), 2-mer (25%), 3-mer (25%) [80 particles per movie]  Unknown: 1-mer (15.38%), 2-mer (14.10%), 3-mer (12.82%), 4-mer (11.54%), 5-mer (10.26%), 6-mer (8.97%), 7-mer (7.69%), 8-mer (6.41%), 9-mer (5.13%), 10-mer (3.85%), 11-mers (2.56% each) and 12-mer (1.28%) [312 particles per movie] |
|  | # frames = 500, # movies = 10, max. photon count = 5, variation in photon count = 20%, lateral sigma = 130 nm, frame size = 1,024 pixels x 1,024 pixels |
|  | Columns EV to EZ (rows 22 to 34) / dataset # 31 copy (2) |
| Figure 4d | Calibration: 1-mer (50%), 2-mer (25%), 3-mer (25%) [80 particles per movie]  Unknown: 1-mer (11.76%), 2-mer (11.03%), 3-mer (10.29%), 4-mer (9.56%), 5-mer (8.82%), 6-mer (8.09%), 7-mer (7.35%), 8-mer (6.62%), 9-mer (5.88%), 10-mer (5.15%), 11-mers (4.41%), 12-mer (3.68%), 13-mer (2.94%), 14-mer (2.21%), 15-mer (1.47%) and 16-mer (0.74%) [544 particles per movie] |
|  | # frames = 500, # movies = 10, max. photon count = 5, variation in photon count = 20%, lateral sigma = 130 nm, frame size = 1,024 pixels x 1,024 pixels |
|  | Columns FH to FL / dataset # 35 |
| Figure 4e | Calibration: 1-mer (50%), 2-mer (25%), 3-mer (25%) [80 particles per movie]  Unknown: 1-mer (9.52%), 2-mer (9.05%), 3-mer (8.57%), 4-mer (8.10%), 5-mer (7.62%), 6-mer (7.14%), 7-mer (6.67%), 8-mer (6.19%), 9-mer (5.71%), 10-mer (5.24%), 11-mers (4.76%), 12-mer (4.29%), 13-mer (3.81%), 14-mer (3.33%), 15-mer (2.86%), 16-mer (2.38%), 17-mer (1.90%), 18-mer (1.43%), 19-mer (0.95%), 20-mer (0.47%) [840 particles per movie] |
|  | # frames = 500, # movies = 10, max. photon count = 5, variation in photon count = 20%, lateral sigma = 130 nm, frame size = 1,024 pixels x 1,024 pixels |
|  | Columns FN to FR / dataset # 36 |
| Figure 4f | Calibration: 1-mer (50%), 2-mer (25%), 3-mer (25%) [80 particles per movie]  Unknown: 1-mer (8%), 2-mer (7.67%), 3-mer (7.33%), 4-mer (7%), 5-mer (6.67%), 6-mer (6.33%), 7-mer (6%), 8-mer (5.67%), 9-mer (5.33%), 10-mer (5%), 11-mers (4.67%), 12-mer (4.33%), 13-mer (4%), 14-mer (3.67%), 15-mer (3.33%), 16-mer (3%), 17-mer (2.67%), 18-mer (2.33%), 19-mer (2%), 20-mer (1.67%), 21-mer (1.33%), 22-mer (1%), 23-mer (0.67%), 24-mer (0.33%) [1200 particles per movie] |
|  | # frames = 500, # movies = 10, max. photon count = 5, variation in photon count = 20%, lateral sigma = 130 nm, frame size = 1,024 pixels x 1,024 pixels |
|  | Columns FT to FX / dataset # 37 |
| Figure 4g | Calibration: 1-mer (50%), 2-mer (25%), 3-mer (25%) [80 particles per movie]  Unknown: 1-mer (28.57%), 3-mer (23.81%), 5-mer (19.10%), 7-mer (14.29%), 9-mer (9.52%) and 11-mer (4.76%) [168 particles per movie] |
|  | # frames = 500, # movies = 10, max. photon count = 5, variation in photon count = 20%, lateral sigma = 130 nm, frame size = 1,024 pixels x 1,024 pixels |
|  | Columns EV to EZ (rows 22 to 34) / dataset # 33 copy (2) |
| Figure 4h | Calibration: 1-mer (50%), 2-mer (25%), 3-mer (25%) [80 particles per movie]  Unknown: 1-mer (22.22%), 3-mer (19.44%), 5-mer (16.67%), 7-mer (13.89%) 9-mer (11.11%), 11-mer (8.33%), 13-mer (5.56%) and 15-mer (2.78%) [288 particles per movie] |
|  | # frames = 500, # movies = 10, max. photon count = 5, variation in photon count = 20%, lateral sigma = 130 nm, frame size = 1,024 pixels x 1,024 pixels |
|  | Columns FZ to GD / dataset # 38 |
| Figure 4i | Calibration: 1-mer (50%), 2-mer (25%), 3-mer (25%) [80 particles per movie]  Unknown: 1-mer (18.18%), 3-mer (16.36%), 5-mer (15.55%), 7-mer (12.73%) 9-mer (10.91%), 11-mer (9.09%), 13-mer (7.27%), 15-mer (5.45%), 17-mer (3.64%) and 19-mer (1.82%) [440 particles per movie] |
|  | # frames = 500, # movies = 10, max. photon count = 5, variation in photon count = 20%, lateral sigma = 130 nm, frame size = 1,024 pixels x 1,024 pixels |
|  | Columns GF to GJ / dataset # 39 |
| Figure 4j | Calibration: 1-mer (50%), 2-mer (25%), 3-mer (25%) [80 particles per movie]  Unknown: 1-mer (15.38%), 3-mer (14.10%), 5-mer (12.82%), 7-mer (11.54%) 9-mer (10.26%), 11-mer (8.97%), 13-mer (7.69%), 15-mer (6.41%), 17-mer (5.13%), 19-mer (3.84%), 21-mer (2.56%) and 23-mer (1.28%) [624 particles per movie] |
|  | # frames = 500, # movies = 10, max. photon count = 5, variation in photon count = 20%, lateral sigma = 130 nm, frame size = 1,024 pixels x 1,024 pixels |
|  | Columns GL to GP / dataset # 40 |
| Figure 4k | Calibration: 1-mer (50%), 2-mer (25%), 3-mer (25%) [80 particles per movie]  Unknown: 1-mer (40%), 4-mer (30%), 7-mer (20%) and 10-mer (10%) [120 particles per movie] |
|  | # frames = 500, # movies = 10, max. photon count = 5, variation in photon count = 20%, lateral sigma = 130 nm, frame size = 1,024 pixels x 1,024 pixels |
|  | Columns GR to GV / dataset # 41 |
| Figure 4l | Calibration: 1-mer (50%), 2-mer (25%), 3-mer (25%) [80 particles per movie]  Unknown: 1-mer (31.37%), 4-mer (25.49%), 7-mer (19.61%), 10-mer (13.73%), 13-mer (7.84%) and 16-mer (1.96%) [204 particles per movie] |
|  | # frames = 500, # movies = 10, max. photon count = 5, variation in photon count = 20%, lateral sigma = 130 nm, frame size = 1,024 pixels x 1,024 pixels |
|  | Columns GX to HB / dataset # 42 |
| Figure 4m | Calibration: 1-mer (50%), 2-mer (25%), 3-mer (25%) [80 particles per movie]  Unknown: 1-mer (25.97%), 4-mer (22.08%), 7-mer (18.18%), 10-mer (14.29%), 13-mer (10.39%), 16-mer (6.49%) and 19-mer (2.60%) [308 particles per movie] |
|  | # frames = 500, # movies = 10, max. photon count = 5, variation in photon count = 20%, lateral sigma = 130 nm, frame size = 1,024 pixels x 1,024 pixels |
|  | Columns HD to HH / dataset # 43 |
| Figure 4n | Calibration: 1-mer (50%), 2-mer (25%), 3-mer (25%) [80 particles per movie]  Unknown: 1-mer (22.22%), 4-mer (19.44%), 7-mer (16.67%), 10-mer (13.89%), 13-mer (11.11%), 16-mer (8.33%), 19-mer (5.56%), 22-mer (2.78%) [432 particles per movie] |
|  | # frames = 500, # movies = 10, max. photon count = 5, variation in photon count = 20%, lateral sigma = 130 nm, frame size = 1,024 pixels x 1,024 pixels |
|  | Columns HJ to HN / dataset # 44 |

**Supplementary Table S1** Information on the stoichiometric configurations and experimental conditions simulated in this study.
