## Supplementary material for "Systematic assessment of the accuracy of subunit counting in biomolecular complexes using automated single molecule brightness analysis": dataAnnotatorAuxFn.m

```
function returnFlag = dataAnnotatorAuxFn(app)
% dataAnnotatorAuxFn- (Auxillary function)
% annotates data using the trained neural network.
%
% Syntax -
% dataAnnotatorAuxFn(app).
%
% Parameters -
% - app: SAS UI class

%% initializing factors
avgFactor = 1;

%% extracting number of files
numFiles = length(app.data.file);

%% looping through files
        n = 1;
for fileId = 1 : numFiles
    
    if strcmp(app.data.file(fileId).type,'Calibration')
        
        %% setting up progress
        app.msgBox.Value = sprintf('%s',['Progress: annotating particles in file ' num2str(fileId) ' out of ' num2str(numFiles)]);
        drawnow;
        
        %% extracting number of particle
        numParticles = length(app.data.file(fileId).particle);
        
        %% validating network
        for particleId = 1 : numParticles
            if strcmp(app.data.file(fileId).particle(particleId).state,'accepted')
                intensity = abs(diff(movmean([app.data.file(fileId).particle(particleId).frame(:).intensity],avgFactor)'));
                gradPeaks = findpeaks((intensity - min(intensity)) ./ (max(intensity) - min(intensity)),'MinPeakHeight',0.5);
                if length(gradPeaks) == 1
                    app.data.file(fileId).particle(particleId).monomeric = true;
                    n = n + 1
                else
                    app.data.file(fileId).particle(particleId).monomeric = false;
                end
            end
        end
    end
end
end
```
