## Supplementary material for "Systematic assessment of the accuracy of subunit counting in biomolecular complexes using automated single molecule brightness analysis": loadParamAuxFn.m

```
function loadParamAuxFn(app)
% loadParamAuxFn() -
% loads SAS parameters.
%
% Syntax -
% exportFn(app).
%
% Parameters -
% - app: SAS UI class

% saving parameters
[importFile,importPath] = uigetfile('*.sp');

% replacing extension with .mat
newFilePath = strrep(fullfile(importPath,importFile),'sp','.mat');

% copying file
copyfile(fullfile(importPath,importFile),newFilePath,'f');

% loading data
try
    paramTemp = load(newFilePath);
catch
    app.msgBox.Value = sprintf('%s',['Error: cannot load data from (' importFile ').']);
    return;
end

% deleting copied file
delete(newFilePath);

% copying paramTemp
app.param = paramTemp.param;

% listing parameters
app.DetectSwitch.Value = app.param.detection.detect;
app.CameraPixelSizeEditField.Value = app.param.detection.pixelSize;
app.CameraOffsetEditField.Value = app.param.detection.cameraOffset;
app.CameraQEEditField.Value = app.param.detection.cameraQE;
app.CameraEMGainEditField.Value = app.param.detection.cameraEMGain;
app.MaximumSigmaEditField.Value = app.param.detection.maxSigma;
app.ROIRadiusEditField.Value = app.param.detection.roiRadius;
app.ProcessSwitch.Value = app.param.procesing.process;
app.AnnotateSwitch.Value = app.param.annotation.annotate;
app.AnalyzeSwitch.Value = app.param.analysis.analyze;
app.LabelingeffeciencyEditField.Value = app.param.analysis.labelingEfficiency;
app.MaximumGMMEditField.Value = app.param.analysis.maxGMM;
app.RefineCheckBox.Value = app.param.analysis.refine;
end
```
