## Supplementary material for "Systematic assessment of the accuracy of subunit counting in biomolecular complexes using automated single molecule brightness analysis": particleLocalizerAuxFn.m

```
function particleLocalizerAuxFn(app,fileId)
% particleLocalizerAuxFn - (Auxillary function)
% localizes particles.
%
% Syntax -
% particleLocalizerAuxFn(app,fileId,particleId)
%
% Parameters -
% - app: SAS UI class.
% - fileId: file #.
% - particleId: particle #.

%% setting up particle image grid for bkgd calculation
roiRadius = 5;
roiWidth = roiRadius * 2 + 1;

%% extracting parameters
cameraPixelSize = app.param.detection.cameraPixelSize;
cameraOffset = app.param.detection.cameraOffset;
cameraEMGain = app.param.detection.cameraEMGain;
cameraQE = app.param.detection.cameraQE;

%% extracting image
image = ((double(app.data.file(fileId).image) - cameraOffset) ./ cameraEMGain) ./ (cameraQE / 100);

%% looping across accepted particles
indices = [];
centroids = [];
numParticles = length(app.data.file(fileId).particle);
for particleId = 1 : numParticles
    if strcmp(app.data.file(fileId).particle(particleId).state,'accepted')
        indices = [indices particleId];
        centroids(particleId,:) = [app.data.file(fileId).particle(particleId).centroid.y ...
            app.data.file(fileId).particle(particleId).centroid.x];
    end
end

%% finding centroids
for particleId = 1 : length(indices)
    subImage(:,:,particleId) = image(round(centroids(indices(particleId),1)) + roiRadius : -1 :...
        round(centroids(indices(particleId),1)) - roiRadius,...
        round(centroids(indices(particleId),2)) - roiRadius:...
        round(centroids(indices(particleId),2)) + roiRadius);
end
maxSubImage = max(subImage(:,:,1 : length(indices)),[],[1,2]);
minSubImage = min(subImage(:,:,1 : length(indices)),[],[1,2]);
fitData(:,1 : length(indices)) = ...
    single(reshape(subImage(:,:,1 : length(indices)),[roiWidth * roiWidth,length(indices)]));
fitInitParams(:,1 : length(indices)) = ...
    single([reshape(maxSubImage(1,1,1:length(indices)),1,[]) - reshape(minSubImage(1,1,1:length(indices)),1,[]) ; ...
    4 * ones(1, length(indices)); ...
    4 * ones(1, length(indices)) ; ...
    1 * ones(1, length(indices)) ; ...
    reshape(minSubImage(1,1,1:length(indices)),1,[])]);

%% fitting particles
[parameters, states, ~, ~, ~] = ...
    cpufit(fitData, ...
    [], ...
    ModelID.GAUSS_2D, ...
    fitInitParams, ...
    [], ...
    [], ...
    [], ...
    []);

%% transferring data to particleData struct
for particleId = 1 : size(fitData,2)
    if states(particleId) == 0
        
        % updating sigma
        app.data.file(fileId).particle(indices(particleId)).sigma = parameters(4,particleId) * cameraPixelSize; 
    else
        app.data.file(fileId).particle(indices(particleId)).state = 'rejected';
    end
end
end
```
