## Supplementary material for "Systematic assessment of the accuracy of subunit counting in biomolecular complexes using automated single molecule brightness analysis": particleRejectorAuxFn.m

```
function particleRejectorAuxFn(app,fileId)
% particleRejectorAuxFn - (Auxillary function)
% rejects particles.
%
% Syntax -
% particleRejectorAuxFn(app,fileId)
%
% Parameters -
% - app: SAS UI class.
% - fileId: file #.
% - particleId: particle #.

%% extracting tolerence and maximum distance
maxSigma = app.param.detection.maxSigma;

%% extracting number of particles
numParticles = length(app.data.file(fileId).particle);

%% looping through particles
for particleId = 1 : numParticles
    
    %% checking if particle is accepted
    if strcmp(app.data.file(fileId).particle(particleId).state,'accepted')
        
        %% extracting sigma
        sigma = app.data.file(fileId).particle(particleId).sigma;
        if sigma > 0 && sigma < maxSigma
        else
            app.data.file(fileId).particle(particleId).state = 'rejected'; 
        end
    end
end
```
