## Supplementary material for "Systematic assessment of the accuracy of subunit counting in biomolecular complexes using automated single molecule brightness analysis": saveParamAuxFn.m

```
function saveParamAuxFn(app)
% saveParamAuxFn() -
% saves SAS parameters.
%
% Syntax -
% exportFn(app).
%
% Parameters -
% - app: SAS UI class

% registering parameters
app.param.detection.detect = app.DetectSwitch.Value;
app.param.detection.pixelSize = app.CameraPixelSizeEditField.Value;
app.param.detection.cameraOffset = app.CameraOffsetEditField.Value;
app.param.detection.cameraQE = app.CameraQEEditField.Value;
app.param.detection.cameraEMGain = app.CameraEMGainEditField.Value;
app.param.detection.maxSigma = app.MaximumSigmaEditField.Value;
app.param.detection.roiRadius = app.ROIRadiusEditField.Value;
app.param.procesing.process = app.ProcessSwitch.Value;
app.param.annotation.annotate = app.AnnotateSwitch.Value;
app.param.analysis.analyze = app.AnalyzeSwitch.Value;
app.param.analysis.labelingEfficiency = app.LabelingeffeciencyEditField.Value;
app.param.analysis.maxGMM = app.MaximumGMMEditField.Value;
app.param.analysis.refine = app.RefineCheckBox.Value;

% saving parameters
param = app.param;
exportPath = uigetdir;
save(fullfile(exportPath,'defparam.sp'),'param');
end
```
