## Supplementary material for "Systematic assessment of the accuracy of subunit counting in biomolecular complexes using automated single molecule brightness analysis": linspecerFnExt.m

```
% function lineStyles = linspecer(N)
% This function creates an Nx3 array of N [R B G] colors
% These can be used to plot lots of lines with distinguishable and nice
% looking colors.
% 
% lineStyles = linspecer(N);  makes N colors for you to use: lineStyles(ii,:)
% 
% colormap(linspecer); set your colormap to have easily distinguishable 
%                      colors and a pleasing aesthetic
% 
% lineStyles = linspecer(N,'qualitative'); forces the colors to all be distinguishable (up to 12)
% lineStyles = linspecer(N,'sequential'); forces the colors to vary along a spectrum 
% 
% % Examples demonstrating the colors.
% 
% LINE COLORS
% N=6;
% X = linspace(0,pi*3,1000); 
% Y = bsxfun(@(x,n)sin(x+2*n*pi/N), X.', 1:N); 
% C = linspecer(N);
% axes('NextPlot','replacechildren', 'ColorOrder',C);
% plot(X,Y,'linewidth',5)
% ylim([-1.1 1.1]);
% 
% SIMPLER LINE COLOR EXAMPLE
% N = 6; X = linspace(0,pi*3,1000);
% C = linspecer(N)
% hold off;
% for ii=1:N
%     Y = sin(X+2*ii*pi/N);
%     plot(X,Y,'color',C(ii,:),'linewidth',3);
%     hold on;
% end
% 
% COLORMAP EXAMPLE
% A = rand(15);
% figure; imagesc(A); % default colormap
% figure; imagesc(A); colormap(linspecer); % linspecer colormap
% 
%   See also NDHIST, NHIST, PLOT, COLORMAP, 43700-cubehelix-colormaps
%%%%%%%%%%%%%%%%%%%%%%%%%%%%%%%%%%%%%%%%%%%%%%%%%%%%%%%%%%%%%%%%%%%%%%%%%%%
% by Jonathan Lansey, March 2009-2013 � Lansey at gmail.com               %
%%%%%%%%%%%%%%%%%%%%%%%%%%%%%%%%%%%%%%%%%%%%%%%%%%%%%%%%%%%%%%%%%%%%%%%%%%%
% 
%% credits and where the function came from
% The colors are largely taken from:
% http://colorbrewer2.org and Cynthia Brewer, Mark Harrower and The Pennsylvania State University
% 
% 
% She studied this from a phsychometric perspective and crafted the colors
% beautifully.
% 
% I made choices from the many there to decide the nicest once for plotting
% lines in Matlab. I also made a small change to one of the colors I
% thought was a bit too bright. In addition some interpolation is going on
% for the sequential line styles.
% 
% 
%%

function lineStyles=linspecerFnExt(N,varargin)

if nargin==0 % return a colormap
    lineStyles = linspecer(128);
    return;
end

if ischar(N)
    lineStyles = linspecer(128,N);
    return;
end

if N<=0 % its empty, nothing else to do here
    lineStyles=[];
    return;
end

% interperet varagin
qualFlag = 0;
colorblindFlag = 0;

if ~isempty(varargin)>0 % you set a parameter?
    switch lower(varargin{1})
        case {'qualitative','qua'}
            if N>12 % go home, you just can't get this.
                warning('qualitiative is not possible for greater than 12 items, please reconsider');
            else
                if N>9
                    warning(['Default may be nicer for ' num2str(N) ' for clearer colors use: whitebg(''black''); ']);
                end
            end
            qualFlag = 1;
        case {'sequential','seq'}
            lineStyles = colorm(N);
            return;
        case {'white','whitefade'}
            lineStyles = whiteFade(N);return;
        case 'red'
            lineStyles = whiteFade(N,'red');return;
        case 'blue'
            lineStyles = whiteFade(N,'blue');return;
        case 'green'
            lineStyles = whiteFade(N,'green');return;
        case {'gray','grey'}
            lineStyles = whiteFade(N,'gray');return;
        case {'colorblind'}
            colorblindFlag = 1;
        otherwise
            warning(['parameter ''' varargin{1} ''' not recognized']);
    end
end      
% *.95
% predefine some colormaps
  set3 = colorBrew2mat({[141, 211, 199];[ 255, 237, 111];[ 190, 186, 218];[ 251, 128, 114];[ 128, 177, 211];[ 253, 180, 98];[ 179, 222, 105];[ 188, 128, 189];[ 217, 217, 217];[ 204, 235, 197];[ 252, 205, 229];[ 255, 255, 179]}');
set1JL = brighten(colorBrew2mat({[228, 26, 28];[ 55, 126, 184]; [ 77, 175, 74];[ 255, 127, 0];[ 255, 237, 111]*.85;[ 166, 86, 40];[ 247, 129, 191];[ 153, 153, 153];[ 152, 78, 163]}'));
set1 = brighten(colorBrew2mat({[ 55, 126, 184]*.85;[228, 26, 28];[ 77, 175, 74];[ 255, 127, 0];[ 152, 78, 163]}),.8);

% colorblindSet = {[215,25,28];[253,174,97];[171,217,233];[44,123,182]};
colorblindSet = {[215,25,28];[253,174,97];[171,217,233]*.8;[44,123,182]*.8};

set3 = dim(set3,.93);

if colorblindFlag
    switch N
        %     sorry about this line folks. kind of legacy here because I used to
        %     use individual 1x3 cells instead of nx3 arrays
        case 4
            lineStyles = colorBrew2mat(colorblindSet);
        otherwise
            colorblindFlag = false;
            warning('sorry unsupported colorblind set for this number, using regular types');
    end
end
if ~colorblindFlag
    switch N
        case 1
            lineStyles = { [  55, 126, 184]/255};
        case {2, 3, 4, 5 }
            lineStyles = set1(1:N);
        case {6 , 7, 8, 9}
            lineStyles = set1JL(1:N)';
        case {10, 11, 12}
            if qualFlag % force qualitative graphs
                lineStyles = set3(1:N)';
            else % 10 is a good number to start with the sequential ones.
                lineStyles = cmap2linspecer(colorm(N));
            end
        otherwise % any old case where I need a quick job done.
            lineStyles = cmap2linspecer(colorm(N));
    end
end
lineStyles = cell2mat(lineStyles);

end

% extra functions
function varIn = colorBrew2mat(varIn)
for ii=1:length(varIn) % just divide by 255
    varIn{ii}=varIn{ii}/255;
end        
end

function varIn = brighten(varIn,varargin) % increase the brightness

if isempty(varargin),
    frac = .9; 
else
    frac = varargin{1}; 
end

for ii=1:length(varIn)
    varIn{ii}=varIn{ii}*frac+(1-frac);
end        
end

function varIn = dim(varIn,f)
    for ii=1:length(varIn)
        varIn{ii} = f*varIn{ii};
    end
end

function vOut = cmap2linspecer(vIn) % changes the format from a double array to a cell array with the right format
vOut = cell(size(vIn,1),1);
for ii=1:size(vIn,1)
    vOut{ii} = vIn(ii,:);
end
end
%%
% colorm returns a colormap which is really good for creating informative
% heatmap style figures.
% No particular color stands out and it doesn't do too badly for colorblind people either.
% It works by interpolating the data from the
% 'spectral' setting on http://colorbrewer2.org/ set to 11 colors
% It is modified a little to make the brightest yellow a little less bright.
function cmap = colorm(varargin)
n = 100;
if ~isempty(varargin)
    n = varargin{1};
end

if n==1
    cmap =  [0.2005    0.5593    0.7380];
    return;
end
if n==2
     cmap =  [0.2005    0.5593    0.7380;
              0.9684    0.4799    0.2723];
          return;
end

frac=.95; % Slight modification from colorbrewer here to make the yellows in the center just a bit darker
cmapp = [158, 1, 66; 213, 62, 79; 244, 109, 67; 253, 174, 97; 254, 224, 139; 255*frac, 255*frac, 191*frac; 230, 245, 152; 171, 221, 164; 102, 194, 165; 50, 136, 189; 94, 79, 162];
x = linspace(1,n,size(cmapp,1));
xi = 1:n;
cmap = zeros(n,3);
for ii=1:3
    cmap(:,ii) = pchip(x,cmapp(:,ii),xi);
end
cmap = flipud(cmap/255);
end

function cmap = whiteFade(varargin)
n = 100;
if nargin>0
    n = varargin{1};
end

thisColor = 'blue';

if nargin>1
    thisColor = varargin{2};
end
switch thisColor
    case {'gray','grey'}
        cmapp = [255,255,255;240,240,240;217,217,217;189,189,189;150,150,150;115,115,115;82,82,82;37,37,37;0,0,0];
    case 'green'
        cmapp = [247,252,245;229,245,224;199,233,192;161,217,155;116,196,118;65,171,93;35,139,69;0,109,44;0,68,27];
    case 'blue'
        cmapp = [247,251,255;222,235,247;198,219,239;158,202,225;107,174,214;66,146,198;33,113,181;8,81,156;8,48,107];
    case 'red'
        cmapp = [255,245,240;254,224,210;252,187,161;252,146,114;251,106,74;239,59,44;203,24,29;165,15,21;103,0,13];
    otherwise
        warning(['sorry your color argument ' thisColor ' was not recognized']);
end

cmap = interpomap(n,cmapp);
end

% Eat a approximate colormap, then interpolate the rest of it up.
function cmap = interpomap(n,cmapp)
    x = linspace(1,n,size(cmapp,1));
    xi = 1:n;
    cmap = zeros(n,3);
    for ii=1:3
        cmap(:,ii) = pchip(x,cmapp(:,ii),xi);
    end
    cmap = (cmap/255); % flipud??
end
```
