## Supplementary material for "Systematic assessment of the accuracy of subunit counting in biomolecular complexes using automated single molecule brightness analysis": gpufit_constrained.m

```
function [parameters, states, chi_squares, n_iterations, time]...
    = gpufit_constrained(data, weights, model_id, initial_parameters, constraints, constraint_types, tolerance, max_n_iterations, parameters_to_fit, estimator_id, user_info)
% Wrapper around the Gpufit mex file.
%
% Optional arguments can be given as empty matrix [].
%
% Default values as specified

%% size checks

% number of input parameter (variable)
if nargin < 12
    user_info = [];
    if nargin < 11
        estimator_id = [];
        if nargin < 10
            parameters_to_fit = [];
            if nargin < 9
                max_n_iterations = [];
                if nargin < 8
                    tolerance = [];
                    if nargin < 7
                        constraint_types = [];
                        if nargin < 6
                            constraints = [];
                            if nargin < 5
                                assert('Not enough parameters');
                            end
                        end
                    end
                end
            end
        end
    end
end

% data is 2D and read number of points and fits
data_size = size(data);
assert(length(data_size) == 2, 'data is not two-dimensional');
n_points = data_size(1);
n_fits = data_size(2);

% consistency with weights (if given)
if ~isempty(weights)
    assert(isequal(data_size, size(weights)), 'Dimension mismatch between data and weights')
end

% initial parameters is 2D and read number of parameters
initial_parameters_size = size(initial_parameters);
assert(length(initial_parameters_size) == 2, 'initial_parameters is not two-dimensional');
n_parameters = initial_parameters_size(1);
assert(n_fits == initial_parameters_size(2), 'Dimension mismatch in number of fits between data and initial_parameters');

% consistency with parameters_to_fit (if given)
if ~isempty(parameters_to_fit)
    assert(size(parameters_to_fit, 1) == n_parameters, 'Dimension mismatch in number of parameters between initial_parameters and parameters_to_fit');
end

%% default values
if isempty(constraints)
    constraints = [];
end

if isempty(constraint_types)
    constraint_types = zeros(n_parameters,'int32');
end

% tolerance
if isempty(tolerance)
    tolerance = 1e-4;
end

% max_n_iterations
if isempty(max_n_iterations)
    max_n_iterations = 25;
end

% estimator_id
if isempty(estimator_id)
    estimator_id = EstimatorID.LSE;
end

% parameters_to_fit
if isempty(parameters_to_fit)
    parameters_to_fit = ones(n_parameters, 1, 'int32');
end

% now only weights and user_info could be not given (empty matrix)

%% type checks

% data, weights (if given), initial_parameters are all single
assert(isa(data, 'single'), 'Type of data is not single');
if ~isempty(weights)
    assert(isa(weights, 'single'), 'Type of weights is not single');
end
assert(isa(initial_parameters, 'single'), 'Type of initial_parameters is not single');

% constraints are either empty or single
if ~isempty(constraints)
    assert(isa(constraints, 'single'), 'Type of constraints is not single');
end

% constraint types are int32
assert(isa(constraint_types,'int32'), 'Type of constraint_types is not int32');

% parameters_to_fit is int32 (cast to int32 if incorrect type)
if ~isa(parameters_to_fit, 'int32')
    parameters_to_fit = int32(parameters_to_fit);
end

% max_n_iterations must be int32 (cast if incorrect type)
if ~isa(max_n_iterations, 'int32')
    max_n_iterations = int32(max_n_iterations);
end

% tolerance must be single (cast if incorrect type)
if ~isa(tolerance, 'single')
    tolerance = single(tolerance);
end

% we don't check type of user_info, but we extract the size in bytes of it
if ~isempty(user_info)
    user_info_info = whos('user_info');
    user_info_size = user_info_info.bytes;
else
    user_info_size = 0;
end


%% run constrained Gpufit taking the time
tic;
[parameters, states, chi_squares, n_iterations] ...
    = GpufitConstrainedMex(...
        data,...
        weights,...
        n_fits,...
        n_points,...
        tolerance,...
        max_n_iterations,...
        estimator_id,...
        initial_parameters,...
        constraints,...
        constraint_types,...
        parameters_to_fit,...
        model_id,...
        n_parameters,...
        user_info,...
        user_info_size);

time = toc;

% reshape the output parameters array to have dimensions
% (n_parameters,n_fits)
parameters = reshape(parameters,n_parameters,n_fits);

end
```
