## Supplementary material for "Systematic assessment of the accuracy of subunit counting in biomolecular complexes using automated single molecule brightness analysis": ModelID.m

```
classdef ModelID
    properties (Constant = true)
        GAUSS_1D = 0
        GAUSS_2D = 1
        GAUSS_2D_ELLIPTIC = 2
        GAUSS_2D_ROTATED = 3
        CAUCHY_2D_ELLIPTIC = 4
        LINEAR_1D = 5
        FLETCHER_POWELL = 6
        BROWN_DENNIS = 7
    end
end
```
