## Supplementary material for "Systematic assessment of the accuracy of subunit counting in biomolecular complexes using automated single molecule brightness analysis": README.gdoc

{
	// WARNING! DO NOT EDIT THIS FILE! ANY CHANGES MADE WILL BE LOST!
	"doc_id" : "1YZDCv7LKmwWT9JBEuziSIFBYGIq2guygJ94rh7Dse1w",
	"resource_key" : "",
	"url" : "https://docs.google.com/open?id=1YZDCv7LKmwWT9JBEuziSIFBYGIq2guygJ94rh7Dse1w"
}
