## Supplementary material for "Systematic assessment of the accuracy of subunit counting in biomolecular complexes using automated single molecule brightness analysis": trackExtFn.m

```
function tracks = trackExtFn(xyzs,maxdisp,param)

%;
% ; see http://glinda.lrsm.upenn.edu/~weeks/idl
% ;   for more information
% ;
% ;+
% ; NAME:
% ; track
% ; PURPOSE:
% ; Constructs n-dimensional trajectories from a scrambled list of
% ; particle coordinates determined at discrete times (e.g. in
% ; consecutive video frames).
% ; CATEGORY:
% ; Image Processing
% ; CALLING SEQUENCE:
% ; result = track( positionlist, maxdisp, param )
% ;  set all keywords in the space below
% ; INPUTS:
% ; positionlist: an array listing the scrambled coordinates and data
% ;     of the different particles at different times, such that:
% ;  positionlist(0:d-1,*): contains the d coordinates and
% ;     data for all the particles, at the different times. must be positve
% ;  positionlist(d,*): contains the time t that the position
% ;     was determined, must be integers (e.g. frame number.  These values must
% ;               be monotonically increasing and uniformly gridded in time.
% ; maxdisp: an estimate of the maximum distance that a particle
% ;     would move in a single time interval.(see Restrictions)
%  OPTIONAL INPUT:
%   param:  a structure containing a few tracking parameters that are
%       needed for many applications.  If param is not included in the
%       function call, then default values are used.  If you set one value
%       make sure you set them all:
% ;         param.mem: this is the number of time steps that a particle can be
% ;             'lost' and then recovered again.  If the particle reappears
% ;             after this number of frames has elapsed, it will be
% ;             tracked as a new particle. The default setting is zero.
% ;             this is useful if particles occasionally 'drop out' of
% ;             the data.
% ;         param.dim: if the user would like to unscramble non-coordinate data
% ;             for the particles (e.g. apparent radius of gyration for
% ;             the particle images), then positionlist should
% ;             contain the position data in positionlist(0:param.dim-1,*)
% ;             and the extra data in positionlist(param.dim:d-1,*). It is then
% ;             necessary to set dim equal to the dimensionality of the
% ;             coordinate data to so that the track knows to ignore the
% ;             non-coordinate data in the construction of the
% ;             trajectories. The default value is two.
% ;         param.good: set this keyword to eliminate all trajectories with
% ;             fewer than param.good valid positions.  This is useful
% ;             for eliminating very short, mostly 'lost' trajectories
% ;             due to blinking 'noise' particles in the data stream.
%;          param.quiet: set this keyword to 1 if you don't want any text
% ; OUTPUTS:
% ; result:  a list containing the original data rows sorted
% ;     into a series of trajectories.  To the original input
% ;     data structure there is appended an additional column
% ;     containing a unique 'id number' for each identified
% ;     particle trajectory.  The result array is sorted so
% ;     rows with corresponding id numbers are in contiguous
% ;     blocks, with the time variable a monotonically
% ;     increasing function inside each block.  For example:
% ;
% ;     For the input data structure (positionlist):
% ;         (x)      (y)      (t)
% ;     pos = 3.60000      5.00000      0.00000
% ;           15.1000      22.6000      0.00000
% ;           4.10000      5.50000      1.00000
% ;           15.9000      20.7000      2.00000
% ;           6.20000      4.30000      2.00000
% ;
% ;     IDL> res = track(pos,5,mem=2)
% ;
% ;     track will return the result 'res'
% ;         (x)      (y)      (t)          (id)
% ;     res = 3.60000      5.00000      0.00000      0.00000
% ;           4.10000      5.50000      1.00000      0.00000
% ;           6.20000      4.30000      2.00000      0.00000
% ;           15.1000      22.6000      0.00000      1.00000
% ;           15.9000      20.7000      2.00000      1.00000
% ;
% ;     NB: for t=1 in the example above, one particle temporarily
% ;     vanished.  As a result, the trajectory id=1 has one time
% ;     missing, i.e. particle loss can cause time gaps to occur
% ;     in the corresponding trajectory list. In contrast:
% ;
% ;     IDL> res = track(pos,5)
% ;
% ;     track will return the result 'res'
% ;         (x)      (y)      (t)          (id)
% ;     res = 15.1000      22.6000      0.00000      0.00000
% ;                   3.60000      5.00000      0.00000      1.00000
% ;               4.10000      5.50000      1.00000      1.00000
% ;               6.20000      4.30000      2.00000      1.00000
% ;               15.9000      20.7000      2.00000      2.00000
% ;
% ;     where the reappeared 'particle' will be labelled as new
% ;     rather than as a continuation of an old particle since
% ;     mem=0.  It is up to the user to decide what setting of
% ;     'mem' will yeild the highest fidelity .
% ;
% ; SIDE EFFECTS:
% ; Produces informational messages.  Can be memory intensive for
% ; extremely large data sets.
% ; RESTRICTIONS:
% ; maxdisp should be set to a value somewhat less than the mean
% ; spacing between the particles. As maxdisp approaches the mean
% ; spacing the runtime will increase significantly. The function
% ; will produce an error message: "Excessive Combinatorics!" if
% ; the run time would be too long, and the user should respond
% ; by re-executing the function with a smaller value of maxdisp.
% ; Obviously, if the particles being tracked are frequently moving
% ; as much as their mean separation in a single time step, this
% ; function will not return acceptable trajectories.
% ; PROCEDURE:
% ; Given the positions for n particles at time t(i), and m possible
% ; new positions at time t(i+1), this function considers all possible
% ; identifications of the n old positions with the m new positions,
% ; and chooses that identification which results in the minimal total
% ; squared displacement. Those identifications which don't associate
% ; a new position within maxdisp of an old position ( particle loss )
% ; penalize the total squared displacement by maxdisp^2. For non-
% ; interacting Brownian particles with the same diffusivity, this
% ; algorithm will produce the most probable set of identifications
% ; ( provided maxdisp >> RMS displacement between frames ).
% ; In practice it works reasonably well for systems with oscillatory,
% ; ballistic, correlated and random hopping motion, so long as single
% ; time step displacements are reasonably small.  NB: multidimensional
% ; functionality is intended to facilitate tracking when additional
% ; information regarding target identity is available (e.g. size or
% ; color).  At present, this information should be rescaled by the
% ; user to have a comparable or smaller (measurement) variance than
% ; the spatial displacements.
% ;
% ; MODIFICATION HISTORY:
% ;  2/93 Written by John C. Crocker, University of Chicago (JFI).
% ;  7/93 JCC fixed bug causing particle loss and improved performance
% ;     for large numbers of (>100) particles.
% ; 11/93 JCC improved speed and memory performance for large
% ;     numbers of (>1000) particles (added subnetwork code).
% ;  3/94 JCC optimized run time for trivial bonds and d<7. (Added
% ;     d-dimensional raster metric code.)
% ;  8/94 JCC added functionality to unscramble non-position data
% ;     along with position data.
% ;  9/94 JCC rewrote subnetwork code and wrote new, more efficient
% ;     permutation code.
% ;  5/95 JCC debugged subnetwork and excessive combinatorics code.
% ; 12/95 JCC added memory keyword, and enabled the tracking of
% ;     newly appeared particles.
% ;  3/96 JCC made inipos a keyword, and disabled the adding of 'new'
% ;     particles when inipos was set.
% ;  3/97 JCC added 'add' keyword, since Chicago users didn't like
% ;     having particle addition be the default.
% ;  9/97 JCC added 'goodenough' keyword to improve memory efficiency
% ;     when using the 'add' keyword and to filter out bad tracks.
% ;       10/97 JCC streamlined data structure to speed runtime for >200
% ;               timesteps.  Changed 'quiet' keyword to 'verbose'. Made
% ;               time labelling more flexible (uniform and sorted is ok).
% ;  9/98 JCC switched trajectory data structure to a 'list' form,
% ;     resolving memory issue for large, noisy datasets.
% ;  2/99 JCC added Eric Weeks's 'uberize' code to post-facto
% ;     rationalize the particle id numbers, removed 'add' keyword.
% ;  1/05 Transmuted to MATLAB by D. Blair
% ;  5/05  ERD Added the param structure to simplify calling.
%    6/05  ERD Added quiet to param structure
%    7/05  DLB Fixed slight bug in trivial bond code
%    3/07  DLB Fixed bug with max disp pointed out by Helene Delanoe-Ayari
%
% ; This code 'track.pro' is copyright 1999, by John C. Crocker.
% ; It should be considered 'freeware'- and may be distributed freely
% ; (outside of the military-industrial complex) in its original form
% ; when properly attributed.
% ;
% ;-

dd = length(xyzs(1,:));

%use default parameters if none given
if nargin==2
    %default values
    memory_b=0; % if mem is not needed set to zero
    goodenough = 0;  % if goodenough is not wanted set to zero
    dim = dd - 1;
    quiet=0;
else
    memory_b    =   param.mem;
    goodenough  =   param.good;
    dim         =   param.dim;
    quiet       =   param.quiet;
end


% checking the input time vector
t = xyzs(:,dd);
st = circshift(t,1);
st = t(2:end) - st(2:end);
if  sum(st(find(st < 0))) ~= 0
    disp('The time vectors is not in order')
    return
end
info = 1;

w = find(st > 0);
z = length(w);
z = z +1;
if isempty(w)
    disp('All positions are at the same time... go back!')
    return
end

% partitioning the data with unique times

%res = unq(t);
% implanting unq directly
indices = find(t ~= circshift(t,-1));
count = length(indices);
if count > 0
    res = indices;
else
    res = length(t) -1;
end
%%%%%%%%%%%%%%%%%%%%%%%

res = [1,res',length(t)];
ngood = res(2) - res(1) + 1;
eyes = 1:ngood;
pos = xyzs(eyes,1:dim);
istart = 2;
n = ngood;

zspan = 50;
if n > 200
    zspan = 20;
end
if n > 500
    zspan = 10;
end
resx = zeros(zspan,n) - 1;

bigresx = zeros(z,n) - 1;
mem = zeros(n,1);
%  whos resx
%  whos bigresx
uniqid = 1:n;
maxid = n;
olist = [0.,0.];

if goodenough > 0
    dumphash = zeros(n,1);
    nvalid = ones(n,1);
end

%  whos eyes;
resx(1,:) = eyes;
% setting up constants
maxdisq = maxdisp^2;

% John calls this the setup for "fancy code" ???
notnsqrd = (sqrt(n*ngood) > 200) & (dim < 7);
notnsqrd = notnsqrd(1);

if notnsqrd
    %;   construct the vertices of a 3x3x3... d-dimensional hypercube
    
    cube = zeros(3^dim,dim);
    
    
    for d=0:dim-1,
        numb = 0;
        for j=0:(3^d):(3^dim)-1,
            cube(j+1:j+(3^(d)),d+1) = numb;
            numb = mod(numb+1,3);
        end
    end
    
    %   calculate a blocksize which may be greater than maxdisp, but which
    %   keeps nblocks reasonably small.
    
    volume = 1;
    for d = 0:dim-1
        minn = min(xyzs(w,d+1));
        maxx = max(xyzs(w,d+1));
        volume = volume * (maxx-minn);
    end
    volume;
    blocksize = max( [maxdisp,((volume)/(20*ngood))^(1.0/dim)] );
end
%   Start the main loop over the frames.
for i=istart:z
    ispan = mod(i-1,zspan)+1;
    %disp(ispan)
    % get new particle positions
    m = res(i+1) - res(i);
    res(i);
    eyes = 1:m;
    eyes = eyes + res(i);
    
    if m > 0
        
        xyi = xyzs(eyes,1:dim);
        found = zeros(m,1);
        
        % THE TRIVIAL BOND CODE BEGINS
        
        if notnsqrd
            %Use the raster metric code to do trivial bonds
            
            % construct "s", a one dimensional parameterization of the space
            % which consists of the d-dimensional raster scan of the volume.)
            
            abi = fix(xyi./blocksize);
            abpos = fix(pos./blocksize);
            si = zeros(m,1);
            spos = zeros(n,1);
            dimm = zeros(dim,1);
            coff = 1.;
            
            for j=1:dim
                minn = min([abi(:,j);abpos(:,j)]);
                maxx = max([abi(:,j);abpos(:,j)]);
                abi(:,j) = abi(:,j) - minn;
                abpos(:,j) = abpos(:,j) - minn;
                dimm(j) = maxx-minn + 1;
                si = si + abi(:,j).*coff;
                spos = spos + abpos(:,j).*coff;
                coff = dimm(j).*coff;
            end
            nblocks = coff;
            % trim down (intersect) the hypercube if its too big to fit in the
            % particle volume. (i.e. if dimm(j) lt 3)
            
            cub = cube;
            deg = find( dimm < 3);
            if ~isempty(deg)
                for j = 0:length(deg)-1
                    cub = cub(find(cub(:,deg(j+1)) < dimm(deg(j+1))),:);
                end
            end
            
            % calculate the "s" coordinates of hypercube (with a corner @ the origin)
            scube = zeros(length(cub(:,1)),1);
            coff = 1;
            for j=1:dim
                scube = scube + cub(:,j).*coff;
                coff = coff*dimm(j);
            end
            
            % shift the hypercube "s" coordinates to be centered around the origin
            
            coff = 1;
            for j=1:dim
                if dimm(j) > 3
                    scube = scube - coff;
                end
                coff = dimm(j).* coff;
            end
            scube = mod((scube + nblocks),nblocks);
            % get the sorting for the particles by their "s" positions.
            [ed,isort] = sort(si);
            
            % make a hash table which will allow us to know which new particles
            % are at a given si.
            strt = zeros(nblocks,1) -1;
            fnsh = zeros(nblocks,1);
            h = find(si == 0);
            lh = length(h);
            if lh > 0
                
                si(h) = 1;
            end
            
            for j=1:m
                if strt(si(isort(j))) == -1
                    strt(si(isort(j))) = j;
                    fnsh(si(isort(j))) = j;
                else
                    fnsh(si(isort(j))) = j;
                end
            end
            if lh > 0
                si(h) = 0;
            end
            coltot = zeros(m,1);
            rowtot = zeros(n,1);
            which1 = zeros(n,1);
            for j=1:n
                
                
                map = fix(-1);
                
                scub_spos = scube + spos(j);
                s = mod(scub_spos,nblocks);
                whzero = find(s == 0 );
                if ~isempty(whzero)
                    nfk = find(s ~=0);
                    s = s(nfk);
                end
                
                w = find(strt(s) ~= -1);
                
                ngood = length(w);
                ltmax=0;
                if ngood ~= 0
                    
                    s = s(w);
                    for k=1:ngood
                        map = [map;isort( strt(s(k)):fnsh(s(k)))];
                    end
                    map = map(2:end);
                    %                     if length(map) == 2
                    %                         if (map(1) - map(2)) == 0
                    %                             map = unique(map);
                    %                          end
                    %                     end
                    %   map = map(umap);
                    %end
                    % find those trival bonds
                    distq = zeros(length(map),1);
                    for d=1:dim
                        distq = distq + (xyi(map,d) - pos(j,d)).^2;
                    end
                    ltmax = distq < maxdisq;
                    
                    rowtot(j) = sum(ltmax);
                    
                    if rowtot(j) >= 1
                        w = find(ltmax == 1);
                        coltot( map(w) ) = coltot( map(w)) +1;
                        which1(j) = map( w(1) );
                    end
                end
                
            end
            
            
            ntrk = fix(n - sum(rowtot == 0));
            
            w = find( rowtot == 1);
            ngood = length(w);
            
            
            if ngood ~= 0
                ww = find(coltot( which1(w) ) == 1);
                ngood = length(ww);
                if ngood ~= 0
                    %disp(size(w(ww)))
                    resx(ispan,w(ww)) = eyes( which1(w(ww)));
                    found( which1( w(ww))) = 1;
                    rowtot( w(ww)) = 0;
                    coltot( which1(w(ww))) = 0;
                end
            end
            
            labely = find( rowtot > 0);
            ngood = length(labely);
            if ngood ~= 0
                labelx = find( coltot > 0);
                
                nontrivial = 1;
            else
                nontrivial = 0;
            end
            
        else
            
            %   or: Use simple N^2 time routine to calculate trivial bonds
            
            % let's try a nice, loopless way!
            % don't bother tracking perm. lost guys.
            wh = find( pos(:,1) >= 0);
            ntrack = length(wh);
            if ntrack == 0
                'There are no valid particles to track idiot!'
                break
            end
            xmat = zeros(ntrack,m);
            count = 0;
            for kk=1:ntrack
                for ll=1:m
                    xmat(kk,ll) = count;
                    count = count+1;
                end
            end
            count = 0;
            for kk=1:m
                for ll=1:ntrack
                    ymat(kk,ll) = count;
                    count = count+1;
                end
            end
            
            xmat = (mod(xmat,m) + 1);
            ymat = (mod(ymat,ntrack) +1)';
            [lenxn,lenxm] = size(xmat);
            %            whos ymat
            %            whos xmat
            %            disp(m)
            
            for d=1:dim
                x = xyi(:,d);
                y = pos(wh,d);
                xm = x(xmat);
                ym = y(ymat(1:lenxn,1:lenxm));
                if size(xm) ~= size(ym)
                    xm = xm';
                end
                
                if d == 1
                    dq = (xm -ym).^2;
                    %dq = (x(xmat)-y(ymat(1:lenxn,1:lenxm))).^2;
                else
                    dq = dq + (xm-ym).^2;
                    %dq = dq + (x(xmat)-y(ymat(1:lenxn,1:lenxm)) ).^2;
                end
            end
            
            ltmax = dq < maxdisq;
            
            % figure out which trivial bonds go with which
            
            rowtot = zeros(n,1);
            rowtot(wh) = sum(ltmax,2);
            
            
            if ntrack > 1
                coltot = sum(ltmax,1);
            else
                coltot = ltmax;
            end
            which1 = zeros(n,1);
            for j=1:ntrack
                [mx, w] = max(ltmax(j,:));
                which1(wh(j)) = w;
            end
            
            ntrk = fix( n - sum(rowtot == 0));
            w= find( rowtot == 1) ;
            ngood = length(w);
            if ngood ~= 0
                ww = find(coltot(which1(w)) == 1);
                ngood = length(ww);
                if ngood ~= 0
                    resx( ispan, w(ww) ) = eyes( which1( w(ww)));
                    found(which1( w(ww))) = 1;
                    rowtot(w(ww)) = 0;
                    coltot(which1(w(ww))) = 0;
                end
            end
            
            labely = find( rowtot > 0);
            ngood = length(labely);
            
            if ngood ~= 0
                labelx = find( coltot > 0);
                nontrivial = 1;
            else
                nontrivial = 0;
            end
        end
        
        %THE TRIVIAL BOND CODE ENDS
        
        if nontrivial
            
            xdim = length(labelx);
            ydim = length(labely);
            
            %  make a list of the non-trivial bonds
            
            bonds = zeros(1,2);
            bondlen = 0;
            
            for j=1:ydim
                distq = zeros(xdim,1);
                
                for d=1:dim
                    %distq
                    distq = distq + (xyi(labelx,d) - pos(labely(j),d)).^2;
                    %distq
                end
                
                w= find(distq <  maxdisq)' - 1;
                ngood = length(w);
                newb = [w;(zeros(1,ngood)+j)];
                
                
                bonds = [bonds;newb'];
                
                bondlen = [ bondlen;distq( w + 1) ];
                
            end
            bonds = bonds(2:end,:);
            
            bondlen = bondlen(2:end);
            numbonds = length(bonds(:,1));
            mbonds = bonds;
            max([xdim,ydim]);
            
            
            if max([xdim,ydim]) < 4
                nclust = 1;
                maxsz = 0;
                mxsz = xdim;
                mysz = ydim;
                bmap = zeros(length(bonds(:,1)+1),1) - 1;
                
            else
                
                
                %   THE SUBNETWORK CODE BEGINS
                lista = zeros(numbonds,1);
                listb = zeros(numbonds,1);
                nclust = 0;
                maxsz = 0;
                thru = xdim;
                
                while thru ~= 0
                    %  the following code extracts connected
                    %   sub-networks of the non-trivial
                    %   bonds.  NB: lista/b can have redundant entries due to
                    %   multiple-connected subnetworks
                    
                    
                    w = find(bonds(:,2) >= 0);
                    %                 size(w)
                    
                    lista(1) = bonds(w(1),2);
                    listb(1) = bonds(w(1),1);
                    bonds(w(1),:) = -(nclust+1);
                    bonds;
                    adda = 1;
                    addb = 1;
                    donea = 0;
                    doneb = 0;
                    if (donea ~= adda) | (doneb ~= addb)
                        true = 0;
                    else
                        true = 1;
                    end
                    
                    while ~true
                        
                        if (donea ~= adda)
                            w = find(bonds(:,2) == lista(donea+1));
                            ngood = length(w);
                            if ngood ~= 0
                                listb(addb+1:addb+ngood,1) = bonds(w,1);
                                bonds(w,:) = -(nclust+1);
                                addb = addb+ngood;
                            end
                            donea = donea+1;
                        end
                        if (doneb ~= addb)
                            w = find(bonds(:,1) == listb(doneb+1));
                            ngood = length(w);
                            if ngood ~= 0
                                lista(adda+1:adda+ngood,1) = bonds(w,2);
                                bonds(w,:) = -(nclust+1);
                                adda = adda+ngood;
                            end
                            doneb = doneb+1;
                        end
                        if (donea ~= adda) | (doneb ~= addb)
                            true = 0;
                        else
                            true = 1;
                        end
                    end
                    
                    [pp,pqx] = sort(listb(1:doneb));
                    %unx =  unq(listb(1:doneb),pqx);
                    %implanting unq directly
                    arr = listb(1:doneb);
                    q = arr(pqx);
                    indices = find(q ~= circshift(q,-1));
                    count = length(indices);
                    if count > 0
                        unx = pqx(indices);
                    else
                        unx = length(q) -1;
                    end
                    %%%%%%%%%%%%%%%%%%%%%%%%%%%%%%%
                    
                    xsz = length(unx);
                    [pp,pqy] = sort(lista(1:donea));
                    %uny =  unq(lista(1:donea),pqy);
                    %implanting unq directly
                    arr = lista(1:donea);
                    q = arr(pqy);
                    indices = find(q ~= circshift(q,-1));
                    count = length(indices);
                    if count > 0
                        uny = pqy(indices);
                    else
                        uny = length(q) -1;
                    end
                    %%%%%%%%%%%%%%%%%%%%%%%%%%%%%%%
                    
                    
                    
                    
                    ysz = length(uny);
                    if xsz*ysz > maxsz
                        maxsz = xsz*ysz;
                        mxsz = xsz;
                        mysz = ysz;
                    end
                    
                    
                    thru = thru -xsz;
                    nclust = nclust + 1;
                end
                bmap = bonds(:,2);
            end
            % THE SUBNETWORK CODE ENDS
            % put verbose in for Jaci
            
            %   THE PERMUTATION CODE BEGINS
            
            for nc =1:nclust
                w = find( bmap == -1*(nc));
                
                nbonds = length(w);
                bonds = mbonds(w,:);
                lensq = bondlen(w);
                [pq,st] = sort( bonds(:,1));
                %un = unq(bonds(:,1),st);
                %implanting unq directly
                arr = bonds(:,1);
                q = arr(st);
                indices = find(q ~= circshift(q,-1));
                count = length(indices);
                if count > 0
                    un = st(indices);
                else
                    un = length(q) -1;
                end
                %%%%%%%%%%%%%%%%%%%%%%%%%%%%%%%%%%%
                
                
                uold = bonds(un,1);
                
                nold = length(uold);
                
                %un = unq(bonds(:,2));
                
                %implanting unq directly
                indices = find(bonds(:,2) ~= circshift(bonds(:,2),-1));
                count = length(indices);
                if count > 0
                    un = indices;
                else
                    un = length(bonds(:,2)) -1;
                end
                %%%%%%%%%%%%%%%%%%%%%%%%%%%%%%%%%%%
                
                unew = bonds(un,2);
                nnew = length(unew);
                
                if nnew > 5
                    rnsteps = 1;
                    for ii =1:nnew
                        rnsteps = rnsteps * length( find(bonds(:,2) == ...
                            unew(ii)));
                        if rnsteps > 5.e+4
                            disp('Warning: difficult combinatorics encountered.')
                        end
                        if rnsteps > 2.e+5
                            disp(['Excessive Combinitorics you FOOL LOOK WHAT YOU HAVE' ...
                                ' DONE TO ME!!!'])
                            return
                        end
                    end
                end
                st = zeros(nnew,1);
                fi = zeros(nnew,1);
                h = zeros(nbonds,1);
                ok = ones(nold,1);
                nlost = (nnew - nold) > 0;
                
                
                for ii=1:nold
                    h(find(bonds(:,1) == uold(ii))) = ii;
                end
                st(1) = 1 ;
                fi(nnew) = nbonds; % check this later
                if nnew > 1
                    sb = bonds(:,2);
                    sbr = circshift(sb,1);
                    sbl = circshift(sb,-1);
                    st(2:end) = find( sb(2:end) ~= sbr(2:end)) + 1;
                    fi(1:nnew-1) = find( sb(1:nbonds-1) ~= sbl(1:nbonds-1));
                end
                %                if i-1 == 13
                %                    hi
                %                end
                checkflag = 0;
                while checkflag ~= 2
                    
                    pt = st -1;
                    lost = zeros(nnew,1);
                    who = 0;
                    losttot = 0;
                    mndisq = nnew*maxdisq;
                    
                    
                    while who ~= -1
                        
                        if pt(who+1) ~= fi(who+1)
                            
                            
                            w = find( ok( h( pt( who+1 )+1:fi( who+1 ) ) ) ); % check this -1
                            ngood = length(w);
                            if ngood > 0
                                if pt(who+1) ~= st(who+1)-1
                                    ok(h(pt(who+1))) = 1;
                                end
                                pt(who+1) = pt(who+1) + w(1);
                                ok(h(pt(who+1))) = 0;
                                if who == nnew -1
                                    ww = find( lost == 0);
                                    dsq = sum(lensq(pt(ww))) + losttot*maxdisq;
                                    
                                    if dsq < mndisq
                                        minbonds = pt(ww);
                                        mndisq = dsq;
                                    end
                                else
                                    who = who+1;
                                end
                            else
                                if ~lost(who+1) & (losttot ~= nlost)
                                    lost(who+1) = 1;
                                    losttot = losttot + 1;
                                    if pt(who+1) ~= st(who+1) -1;
                                        ok(h(pt(who+1))) = 1;
                                    end
                                    if who == nnew-1
                                        ww = find( lost == 0);
                                        dsq = sum(lensq(pt(ww))) + losttot*maxdisq;
                                        if dsq < mndisq
                                            minbonds = pt(ww);
                                            mndisq = dsq;
                                        end
                                    else
                                        who = who + 1;
                                    end
                                    
                                else
                                    if pt(who+1) ~= (st(who+1) -1)
                                        ok(h(pt(who+1))) = 1;
                                    end
                                    pt(who+1) = st(who+1) -1;
                                    if lost(who+1)
                                        lost(who+1) = 0;
                                        losttot = losttot -1;
                                    end
                                    who = who -1;
                                end
                            end
                        else
                            if ~lost(who+1) & (losttot ~= nlost)
                                lost(who+1) = 1;
                                losttot = losttot + 1;
                                if pt(who+1) ~= st(who+1)-1
                                    ok(h(pt(who+1))) = 1;
                                end
                                if who == nnew -1
                                    ww = find( lost == 0);
                                    dsq = sum(lensq(pt(ww))) + losttot*maxdisq;
                                    
                                    if dsq < mndisq
                                        minbonds = pt(ww);
                                        mndisq = dsq;
                                    end
                                else
                                    who = who + 1;
                                end
                            else
                                if pt(who+1) ~= st(who+1) -1
                                    ok(h(pt(who+1))) = 1;
                                end
                                pt(who+1) = st(who+1) -1;
                                if lost(who+1)
                                    lost(who+1) = 0;
                                    losttot = losttot -1;
                                end
                                who = who -1;
                            end
                        end
                    end
                    
                    checkflag = checkflag + 1;
                    if checkflag == 1
                        plost = min([fix(mndisq/maxdisq) , (nnew -1)]);
                        if plost > nlost
                            nlost = plost;
                        else
                            checkflag = 2;
                        end
                    end
                    
                end
                %   update resx using the minimum bond configuration
                
                resx(ispan,labely(bonds(minbonds,2))) = eyes(labelx(bonds(minbonds,1)+1));
                found(labelx(bonds(minbonds,1)+1)) = 1;
                
            end
            
            %   THE PERMUTATION CODE ENDS
        end
        
        w = find(resx(ispan,:) >= 0);
        nww = length(w);
        
        if nww > 0
            pos(w,:) = xyzs( resx(ispan,w) , 1:dim);
            if goodenough > 0
                nvalid(w) = nvalid(w) + 1;
            end
        end  %go back and add goodenough keyword thing
        newguys = find(found == 0);
        
        nnew = length(newguys);
        
        if (nnew > 0) % & another keyword to workout inipos
            newarr = zeros(zspan,nnew) -1;
            resx = [resx,newarr];
            
            resx(ispan,n+1:end) = eyes(newguys);
            pos = [[pos];[xyzs(eyes(newguys),1:dim)]];
            nmem = zeros(nnew,1);
            mem = [mem;nmem];
            nun = 1:nnew;
            uniqid = [uniqid,((nun) + maxid)];
            maxid = maxid + nnew;
            if goodenough > 0
                dumphash = [dumphash;zeros(1,nnew)'];
                nvalid = [nvalid;zeros(1,nnew)'+1];
            end
            % put in goodenough
            n = n + nnew;
            
        end
        
    else
        ' Warning- No positions found for t='
    end
    w = find( resx(ispan,:) ~= -1);
    nok = length(w);
    if nok ~= 0
        mem(w) =0;
    end
    
    mem = mem + (resx(ispan,:)' == -1);
    wlost = find(mem == memory_b+1);
    nlost =length(wlost);
    
    if nlost > 0
        pos(wlost,:) = -maxdisp;
        if goodenough > 0
            wdump = find(nvalid(wlost) < goodenough);
            ndump = length(wdump);
            if ndump > 0
                dumphash(wlost(wdump)) = 1;
            end
        end
        % put in goodenough keyword stuff if
    end
    if (ispan == zspan) | (i == z)
        nold = length(bigresx(1,:));
        nnew = n-nold;
        if nnew > 0
            newarr = zeros(z,nnew) -1;
            bigresx = [bigresx,newarr];
        end
        if goodenough > 0
            if (sum(dumphash)) > 0
                wkeep = find(dumphash == 0);
                nkeep = length(wkeep);
                resx = resx(:,wkeep);
                bigresx = bigresx(:,wkeep);
                pos = pos(wkeep,:);
                mem = mem(wkeep);
                uniqid = uniqid(wkeep);
                nvalid = nvalid(wkeep);
                n = nkeep;
                dumphash = zeros(nkeep,1);
            end
        end
        
        % again goodenough keyword
        if quiet~=1
            disp(strcat(num2str(i), ' of ' ,num2str(z), ' done.  Tracking  ',num2str(ntrk),' particles  ', num2str(n),' tracks total'));
        end
        bigresx(i-(ispan)+1:i,:) = resx(1:ispan,:);
        resx = zeros(zspan,n) - 1;
        
        
        wpull = find(pos(:,1) == -maxdisp);
        npull = length(wpull);
        
        if npull > 0
            lillist = zeros(1,2);
            for ipull=1:npull
                wpull2 = find(bigresx(:,wpull(ipull)) ~= -1);
                npull2 = length(wpull2);
                thing = [bigresx(wpull2,wpull(ipull)),zeros(npull2,1)+uniqid(wpull(ipull))];
                lillist = [lillist;thing];
                
            end
            olist = [[olist];[lillist(2:end,:)]];
            
        end
        
        
        
        wkeep = find(pos(:,1) >= 0);
        nkeep = length(wkeep);
        if nkeep == 0
            'Were going to crash now, no particles....'
        end
        resx = resx(:,wkeep);
        bigresx = bigresx(:,wkeep);
        pos = pos(wkeep,:);
        mem = mem(wkeep);
        uniqid = uniqid(wkeep);
        n = nkeep;
        dumphash = zeros(nkeep,1);
        if goodenough > 0
            nvalid = nvalid(wkeep);
        end
    end
    
end

if goodenough > 0
    nvalid = sum(bigresx >= 0 ,1);
    wkeep = find(nvalid >= goodenough);
    nkeep = length(wkeep);
    if nkeep == 0
        for i=1:10
            disp('You are not going any further, check your params and data')
        end
        disp('the code broke at line 1045')
        return
    end
    if nkeep < n
        bigresx = bigresx(:,wkeep);
        n = nkeep;
        uniqid = uniqid(wkeep);
        pos = pos(wkeep,:);
    end
end


wpull = find( pos(:,1) ~= -2*maxdisp);
npull = length(wpull);
if npull > 0
    lillist = zeros(1,2);
    for ipull=1:npull
        wpull2 = find(bigresx(:,wpull(ipull)) ~= -1);
        npull2 = length(wpull2);
        thing = [bigresx(wpull2,wpull(ipull)),zeros(npull2,1)+uniqid(wpull(ipull))];
        lillist = [lillist;thing];
    end
    olist = [olist;lillist(2:end,:)];
end

olist = olist(2:end,:);
%bigresx = 0;
%resx = 0;

nolist = length(olist(:,1));
res = zeros(nolist,dd+1);
for j=1:dd
    res(:,j) = xyzs(olist(:,1),j);
end
res(:,dd+1) = olist(:,2);

% this is uberize included for simplicity of a single monolithic code

ndat=length(res(1,:));
newtracks=res;


%u=unq(newtracks(:,ndat));

% inserting unq
indices = find(newtracks(:,ndat) ~= circshift(newtracks(:,ndat),-1));
count = length(indices);
if count > 0
    u = indices;
else
    u = length(newtracks(:,ndat)) -1;
end


ntracks=length(u);
u=[0;u];
for i=2:ntracks+1
    newtracks(u(i-1)+1:u(i),ndat) = i-1;
end

% end of uberize code

tracks = newtracks;
end
```
