## Supplementary material for "Systematic assessment of the accuracy of subunit counting in biomolecular complexes using automated single molecule brightness analysis": analyzeFn.m

```
function returnFlag = analyzeFn(app)
% analyzeFn() -
% analyzes data processed by SAS.
%
% Syntax -
% analyzeFn(app).
%
% Parameters -
% - app: SAS UI class

%% initializing returnFlag
returnFlag = false;

%% displaying SAS progress
app.msgBox.Value = sprintf('%s','Progress: data analysis started.');
drawnow;

%% calling ParticleDetector
app.msgBox.Value = sprintf('%s','Progress: generating and fitting kernel estimator.');
drawnow;
KernelGeneratorAndFitter(app);

%% calling LabelingCorrector
app.msgBox.Value = sprintf('%s','Progress: correcting for labelling.');
drawnow;
LabellingCorrector(app);

%% exporting files relevant to this function
app.msgBox.Value = sprintf('%s','Progress: exporting data.');
drawnow;
SpecificExport(app);

app.msgBox.Value = sprintf('%s','Done.');
drawnow;
end

%%====================KernelGenerator=====================%%
function KernelGeneratorAndFitter(app)
global x0 s refineId refinedIndex numGaussCalib meanCalib numGaussCurrent;

% extracting number of files
numFiles = length(app.data.file);

% extracting maximum number of Gaussian mixture
maxGMM = app.param.analysis.maxGMM;

% initializing intensity vector
intCalibration = [];
intUnknown = [];

% looping over files
for fileId = 1 : numFiles
    
    % checking if file is a calibration file
    if strcmp(app.data.file(fileId).type,'Calibration')
        
        % extracting number of particles
        numParticles = length(app.data.file(fileId).particle);
        
        % looping over particles
        for particleId = 1 : numParticles
            
            % checking if particle is accepted
            if strcmp(app.data.file(fileId).particle(particleId).state,'accepted')
                
                if app.data.file(fileId).particle(particleId).monomeric

                    % calculating the intensity
                    intCalibration = [intCalibration ...
                        app.data.file(fileId).particle(particleId).maxIntensity - ...
                        app.data.file(fileId).particle(particleId).minIntensity];
                end
            end
        end
    else
        
        % extracting number of particles
        numParticles = length(app.data.file(fileId).particle);
        
        % looping over particles
        for particleId = 1 : numParticles
            
            % checking if particle is accepted
            if strcmp(app.data.file(fileId).particle(particleId).state,'accepted')

                % calculating the intensity
                intUnknown = [intUnknown ...
                    app.data.file(fileId).particle(particleId).maxIntensity - ...
                    app.data.file(fileId).particle(particleId).minIntensity];
            end
        end
    end
end

% fitting calibration data
numGaussCalib = 2;
x_calib = min(intCalibration) : 1 : max(intCalibration);
pd = fitdist(intCalibration','Kernel','BandWidth',5);
y_calib = pdf(pd,x_calib);
[~,peakLoc] = findpeaks((y_calib - min(y_calib)) ./ (max(y_calib) - min(y_calib)),'MinPeakHeight',0.4);
param = lsqcurvefit(@GMM,...
    [y_calib(peakLoc(1)) * ones(1,numGaussCalib) x_calib(peakLoc(1)) x_calib(peakLoc(1)) / 4 x_calib(peakLoc(1)) / 4],...
    x_calib,...
    y_calib,...
    [0 0 x_calib(peakLoc(1)) 0 0],...
    [inf inf x_calib(peakLoc(1)) inf inf]);
app.data.curve.y_calib = y_calib;
app.data.curve.x_calib = x_calib;
app.data.curve.param = param;

% finding mean and sigma of the monomeric species
meanCalibration = param(numGaussCalib + 1);
sigmaCalibration = param(numGaussCalib + 2);

% constructing maximum number of Gaussians
numGauss = floor((meanCalibration / sigmaCalibration) ^ 2);
numGauss = min([numGauss maxGMM]);

% constructing mean and std vector
x0 = [];
s = [];
if app.param.analysis.refine
    numRefines = 20;
else
    numRefines = 1;
end
for refineId = 1 : numRefines
    for gaussId = 1 : numGauss
        x0(refineId,gaussId) = gaussId * (meanCalibration - ceil(numRefines/2) + refineId);
        s(refineId,gaussId) = sqrt(gaussId) * sigmaCalibration;
    end
end

% constructing a kernel distribution
x_unknown = min(intUnknown) : 1 : max(intUnknown);
pd = fitdist(intUnknown','Kernel','BandWidth',5);
y_unknown = pdf(pd,x_unknown);
app.data.curve.y_unknown = y_unknown;
app.data.curve.x_unknown = x_unknown;

% fitting gaussian mixture model to data
for refineId = 1 : numRefines
    minResidual = inf;
    for gaussId = 1 : numGauss
        numGaussCurrent = gaussId;
        [species,~,residual,~,~] = lsqcurvefit(@ModelGenerator,double((1 / gaussId) .* ones(1,gaussId)),x_unknown,y_unknown,...
            zeros(1,gaussId),inf * ones(1,gaussId));
        if sqrt(sum(residual .^ 2)) < 0.95 * minResidual
            minResidual = sqrt(sum(residual .^ 2));
            chosenSpecies = species;
        end
    end
    refinedResidual(refineId) = minResidual;
    refinedSpecies.refine{refineId} = chosenSpecies;
end

% choosing species after refinement
[~,refinedIndex] = min(refinedResidual);
if (refinedIndex == 1 || refinedIndex == numRefines) && app.param.analysis.refine
     refinedIndex = 5;
end
app.data.species.raw = refinedSpecies.refine{refinedIndex};
app.data.curve.refinedParam.x0 = x0(refinedIndex,:);
app.data.curve.refinedParam.s = s(refinedIndex,:);
end

%%====================KernelGenerator=====================%%
function LabellingCorrector(app)

% expressing propotions as percentages
app.data.species.before = (100 .* app.data.species.raw) ./ sum(app.data.species.raw);

% constructing matrix with binomial probabilities
dim = numel(app.data.species.before);

% constructing binomial distribution
for n = 1 : dim
    for x = 1 : dim
        binomialPdf(x,n) = binopdf(x,n,app.param.analysis.labelingEfficiency / 100);
    end
end

equalityConstraint_A = ones(1,dim);
equalityConstraint_B = 100;
lowerBound = zeros(dim,1);
upperBound = 100 .* ones(dim,1);

% calculating number of species
[species,~,~,~,~,~] = lsqlin(binomialPdf,app.data.species.before,[],[],equalityConstraint_A,equalityConstraint_B,lowerBound,upperBound);
app.data.species.after = 100 .* species ./ sum(species);
end

%%====================KernelGenerator=====================%%
function SpecificExport(app)
global s x0 numGaussCalib refinedIndex

% creating new folder
mkdir(fullfile(app.param.paths.calibrationAndUnknownData,'analysis'));

% calculating file handle and opening file
fileHandle = fopen(fullfile(app.param.paths.calibrationAndUnknownData,...
    'analysis',...
    'Unknown_species proportion(summary).txt'),'w');

% writing to text file proportions before correcting for labelling efficiency      
fprintf(fileHandle,'%s\n','Proportions before correcting for labelling efficiency');
for unitId = 1 : length(app.data.species.before)
     fprintf(fileHandle,'%d\t%d\n',unitId,app.data.species.before(unitId));
end

% line skipping
fprintf(fileHandle,'\n');

% writing to text file proportions after correcting for labelling efficiency      
fprintf(fileHandle,'%s\n','Proportions after correcting for labelling efficiency');
for unitId = 1 : length(app.data.species.after)
     fprintf(fileHandle,'%d\t%d\n',unitId,app.data.species.after(unitId));
end

% closing file
fclose(fileHandle);

% creating figure (bar graph of species before labelling correction)
figHandle = figure('visible','off');

% drawing bar graph
bar(1 : length(app.data.species.before),app.data.species.before);
xlabel('Species');
ylabel('Proportions (%)')

% saving figure
saveas(figHandle,fullfile(app.param.paths.calibrationAndUnknownData,...
    'analysis',...
    'Unknown_species proportion(before labelling correction).png'));

% creating figure (bar graph of species after labelling correction)
figHandle = figure('visible','off');

% drawing bar graph
bar(1 : length(app.data.species.after),app.data.species.after);
xlabel('Species');
ylabel('Proportions (%)')

% saving figure
saveas(figHandle,fullfile(app.param.paths.calibrationAndUnknownData,...
    'analysis',...
    'Unknown_species proportion(after labelling correction).png'));

% saving PDF of unknown data in text file
% calculating file handle and opening file
fileHandle = fopen(fullfile(app.param.paths.calibrationAndUnknownData,...
    'analysis',...
    'Unknown_PDF.txt'),'w'); 

% creating figure (bar graph of species after labelling correction)
figHandle = figure('visible','off');

% drawing kernel density estimator
plot(app.data.curve.x_unknown,app.data.curve.y_unknown);
hold on;

% recording kernel density estimator in text file
fprintf(fileHandle,'%s\n','Kernel Density Estimator');
fprintf(fileHandle,'%s\t%s\n','Intensity (Photons)','Frequency');
for indexId = 1 : length(app.data.curve.x_unknown)
    fprintf(fileHandle,'%f\t%f\n',...
        app.data.curve.x_unknown(indexId),...
        app.data.curve.y_unknown(indexId));
end
fprintf(fileHandle,'\n');

% drawing gaussian mixture
for gaussId = 1 : length(app.data.species.before)
    y = [];
    for x = app.data.curve.x_unknown
       y = [y (app.data.species.raw(gaussId) ./ ...
           (s(refinedIndex,gaussId) .* sqrt(2 * pi))) .* exp(-((x - x0(refinedIndex,gaussId)) .^ 2) ./ (2 .* (s(refinedIndex,gaussId) ^ 2)))]; 
    end
    plot(app.data.curve.x_unknown,y);
    hold on;
    fprintf(fileHandle,'%s\t%d\n','Gaussian Curve',gaussId);
    fprintf(fileHandle,'%s\t%s\n','Intensity (Photons)','Frequency');
    for indexId = 1 : length(app.data.curve.x_unknown)
        fprintf(fileHandle,'%f\t%f\n',...
            app.data.curve.x_unknown(indexId),...
            y(indexId));
    end
    fprintf(fileHandle,'\n');
end
hold off;
xlabel('Intensity (photons)');
ylabel('Frequency')

% saving figure
saveas(figHandle,fullfile(app.param.paths.calibrationAndUnknownData,...
    'analysis',...
    'Unknown_PDF.png'));

% closing file
fclose(fileHandle);

% saving PDF of calibration data in text file
% calculating file handle and opening file
fileHandle = fopen(fullfile(app.param.paths.calibrationAndUnknownData,...
    'analysis',...
    'Calibration_PDF.txt'),'w'); 

% creating figure (bar graph of species after labelling correction)
figHandle = figure('visible','off');

% drawing kernel density estimator
plot(app.data.curve.x_calib,app.data.curve.y_calib);
hold on;

% recording kernel density estimator in text file
fprintf(fileHandle,'%s\n','Kernel Density Estimator');
fprintf(fileHandle,'%s\t%s\n','Intensity (Photons)','Frequency');
for indexId = 1 : length(app.data.curve.x_calib)
    fprintf(fileHandle,'%f\t%f\n',...
        app.data.curve.x_calib(indexId),...
        app.data.curve.y_calib(indexId));
end
fprintf(fileHandle,'\n');

% drawing gaussian mixture
for gaussId = 1 : numGaussCalib
    y = [];
    for x = app.data.curve.x_calib
       y = [y (app.data.curve.param(gaussId) .* exp(-((x - (app.data.curve.param(numGaussCalib + 1) * gaussId)) .^ 2) ./ (2 .* ((app.data.curve.param(numGaussCalib + 1 + gaussId) * sqrt(gaussId)) ^ 2))))]; 
    end
    plot(app.data.curve.x_calib,y);
    hold on;
    fprintf(fileHandle,'%s\t%d\n','Gaussian Curve',gaussId);
    fprintf(fileHandle,'%s\t%s\n','Intensity (Photons)','Frequency');
    for indexId = 1 : length(app.data.curve.x_calib)
        fprintf(fileHandle,'%f\t%f\n',...
            app.data.curve.x_calib(indexId),...
            y(indexId));
    end
    fprintf(fileHandle,'\n');
end
hold off;
xlabel('Intensity (photons)');
ylabel('Frequency')

% saving figure
saveas(figHandle,fullfile(app.param.paths.calibrationAndUnknownData,...
    'analysis',...
    'Calibration_PDF.png'));

% closing file
fclose(fileHandle);

% saving coordinates and intensities of calibration and unknown data in text file
% calculating file handle and opening file
fileHandle = fopen(fullfile(app.param.paths.calibrationAndUnknownData,...
    'analysis',...
    'Calibration_raw.txt'),'w'); 
fprintf(fileHandle,'%s\t%s\t%s\n','Intensity','x','y');
numFiles = length(app.data.file);
for fileId = 1 : numFiles
    
    % checking if file is a calibration file
    if strcmp(app.data.file(fileId).type,'Calibration')
        
        % extracting number of particles
        numParticles = length(app.data.file(fileId).particle);
        
        % looping over particles
        for particleId = 1 : numParticles
            
            % checking if particle is accepted
            if strcmp(app.data.file(fileId).particle(particleId).state,'accepted')
                
                if app.data.file(fileId).particle(particleId).monomeric
                    
                    % printing to file
                    fprintf(fileHandle,'%f\t%f\t%f\n',...
                        app.data.file(fileId).particle(particleId).maxIntensity - ...
                        app.data.file(fileId).particle(particleId).minIntensity,...
                        app.data.file(fileId).particle(particleId).centroid.x,...
                        app.data.file(fileId).particle(particleId).centroid.y);
                end
            end
        end
    end
end

% closing file
fclose(fileHandle);

fileHandle = fopen(fullfile(app.param.paths.calibrationAndUnknownData,...
    'analysis',...
    'Unknown_raw.txt'),'w'); 
fprintf(fileHandle,'%s\t%s\t%s\n','Intensity','x','y');
numFiles = length(app.data.file);
for fileId = 1 : numFiles
    
    % checking if file is a calibration file
    if strcmp(app.data.file(fileId).type,'Unknown')
        
        % extracting number of particles
        numParticles = length(app.data.file(fileId).particle);
        
        % looping over particles
        for particleId = 1 : numParticles
            
            % checking if particle is accepted
            if strcmp(app.data.file(fileId).particle(particleId).state,'accepted')
                 
                    % printing to file
                    fprintf(fileHandle,'%f\t%f\t%f\n',...
                        app.data.file(fileId).particle(particleId).maxIntensity - ...
                        app.data.file(fileId).particle(particleId).minIntensity,...
                        app.data.file(fileId).particle(particleId).centroid.x,...
                        app.data.file(fileId).particle(particleId).centroid.y);
            end
        end
    end
end

% closing file
fclose(fileHandle);

% saving all data
data = app.data;
save(fullfile(app.param.paths.calibrationAndUnknownData,'All data.mat'),'data');
end

%%====================ModelGenerator=====================%%
function y = ModelGenerator(p,x)
global x0 s refineId numGaussCurrent
y = zeros(size(x));
for xId = 1 : length(x)
    for gaussId = 1 : numGaussCurrent
        y(xId) = y(xId) + (p(gaussId) ./ (s(refineId,gaussId) .* sqrt(2 * pi))) .* exp(-((x(xId) - x0(refineId,gaussId)) .^ 2) ./ (2 .* (s(refineId,gaussId) ^ 2)));
    end
end
end

%%====================FunctionGenerator=====================%%
function y = GMM(param,x)
global numGaussCalib
y = zeros(size(x));
for xId = 1 : length(x)
    for gaussId = 1 : numGaussCalib
        y(xId) = y(xId) + (param(gaussId) .* exp(-((x(xId) - (param(numGaussCalib + 1) * gaussId)) .^ 2) ./ (2 .* ((param(numGaussCalib + 1 + gaussId) * sqrt(gaussId)) ^ 2))));
    end
end
end
```
