## Supplementary material for "Systematic assessment of the accuracy of subunit counting in biomolecular complexes using automated single molecule brightness analysis": annotateFn.m

```
function returnFlag = annotateFn(app)
% annotateFn() - 
% annotates calibration data saved by SAS.
%
% Syntax - 
% annotateFn(app).
%
% Parameters -
% - app: SAS UI class

%% initializing returnFlag
returnFlag = false;

%% issuing initial error statements
if isempty(app.param.paths.calibrationAndUnknownData)
    returnFlag = true;
    app.msgBox.Value = sprintf('%s','Error: no data path selected.');
    return;
end

%% checking availability of calibration data
dataFileList = dir(fullfile(app.param.paths.calibrationAndUnknownData,'Calibration','native','*.sd'));
if isempty(dataFileList)
    returnFlag = true;
    app.msgBox.Value = sprintf('%s','Error: no calibration data in selected path.');
    return;
end

%% calling data annotator
DataAnnotator(app);

%% calling exportFn
exportFn(app);
end

%%====================DataAnnotator=====================%%
function DataAnnotator(app)

% calling dataAnnotatorAuxFn
dataAnnotatorAuxFn(app);
end
```
