## Supplementary material for "Systematic assessment of the accuracy of subunit counting in biomolecular complexes using automated single molecule brightness analysis": detectFn.m

```
function returnFlag = detectFn(app)
% detectFn() -
% detect diffraction-limited blobs in Field of View
%
% Syntax -
% detectFn(app).
%
% Parameters -
% - app: SAS UI class

%% initializing returnFlag
returnFlag = false;

%% issuing initial error statements
if isempty(app.param.paths.calibrationAndUnknownData)
    returnFlag = true;
    app.msgBox.Value = sprintf('%s','Error: no data path selected.');
    return;
end

%% checking availability of calibration and unknown data
cd(fullfile(app.param.paths.calibrationAndUnknownData,'Calibration'));
dataFileList = dir('*.tif');
if isempty(dataFileList)
    returnFlag = true;
    app.msgBox.Value = sprintf('%s','Error: no calibration data in selected patb.');
    return;
end
cd(fullfile(app.param.paths.calibrationAndUnknownData,'Unknown'));
dataFileList = dir('*.tif');
if isempty(dataFileList)
    returnFlag = true;
    app.msgBox.Value = sprintf('%s','Error: no unknown data in selected patb.');
    return;
end

%% displaying progress
app.msgBox.Value = sprintf('%s','Progress: detection started.');

%% looping though both folders, calibrtion (1) and unknown data(2)
for roundId = 1 : 2
    
    %% initializing struct
    app.data = struct();
    
    %% reading input files
    if roundId == 1
        cd(fullfile(app.param.paths.calibrationAndUnknownData,'Calibration'));
    else
        cd(fullfile(app.param.paths.calibrationAndUnknownData,'Unknown'));
    end
    inputFiles = dir('*.tif');
    
    %% reading number of files
    numFiles = numel(inputFiles);
    
    %% looping through files
    for fileId = 1 : numFiles
        
        % registering state
        app.data.file(fileId).state = 'detected';
        
        % registering type
        if roundId == 1
            app.data.file(fileId).type = 'Calibration';
        else
            app.data.file(fileId).type = 'Unknown';
        end
        
        %% extracting file path
        if numFiles == 1
            fileName = inputFiles.name;
            fileFolder = inputFiles.folder;
        else
            fileName = inputFiles(fileId).name;
            fileFolder = inputFiles(fileId).folder;
        end
        filePath = fullfile(fileFolder,fileName);
        app.data.file(fileId).name = fileName;
        
        %% reading first frame
        try
            image = imread(filePath,1);
            for frameId = 2 : 5
                image = image + imread(filePath,frameId);
            end
            image = image ./ 5;
        catch
            returnFlag = true;
            app.msgBox.Value = sprintf('%s',['Error: cannot read image file (' fileName ').']);
            return;
        end
        
        %% adding image to particleData struct
        app.data.file(fileId).image = double(image);
        
        %% setting up progress
        app.msgBox.Value = sprintf('%s',['Progress: detecting particles in file ' num2str(fileId) ' out of ' num2str(numFiles) '. Classifying regional maximas.']);
        drawnow;
        
        %% calling ParticleDetector
        ParticleDetector(app,fileId);
        
        %% setting up progress
        app.msgBox.Value = sprintf('%s',['Progress: detecting particles in file ' num2str(fileId) ' out of ' num2str(numFiles) '. Localizing particles.']);
        drawnow;
        
        ParticleLocalizer(app,fileId);
        
        %% setting up progress
        app.msgBox.Value = sprintf('%s',['Progress: detecting particles in file ' num2str(fileId) ' out of ' num2str(numFiles) '. Rejecting non-singular particles.']);
        drawnow;
        
        ParticleRejector(app,fileId);
    end
    
    %% exporting generic .m files
    exportFn(app);
    
    %% exporting files relevant to this function
    SpecificExport(app);
end

%% displaying progress
app.msgBox.Value = sprintf('%s','Progress: detection complete.');
end

%%====================ParticleDetector=====================%%
function ParticleDetector(app,fileId)

% extracting parameters
cameraOffset = app.param.detection.cameraOffset;
cameraEMGain = app.param.detection.cameraEMGain;
cameraQE = app.param.detection.cameraQE;
roiRadius = 5;
roiWidth = (roiRadius * 2) + 1;

% extracting image
imageRaw = ((double(app.data.file(fileId).image) - cameraOffset) ./ cameraEMGain) ./ (cameraQE / 100);

% thresholding image
thresholdImage = imregionalmax(imageRaw,8);

% extracting connected objects from threshold image
[centroidTemp_y,centroidTemp_x] = ind2sub(size(thresholdImage),find(thresholdImage == true));
centroids = [centroidTemp_x centroidTemp_y];

% creating an array of all particles in an image
imageVec = zeros(roiWidth,roiWidth,1,size(centroids,1));
for particleId = 1 : size(centroids,1)
    try
        imageVec(:,:,1,particleId) = imageRaw(round(centroids(particleId,2)) - roiRadius :...
            round(centroids(particleId,2)) + roiRadius,...
            round(centroids(particleId,1)) - roiRadius:...
            round(centroids(particleId,1)) + roiRadius);
        imageVec(:,:,:,particleId) = (imageVec(:,:,:,particleId) - min(min(imageVec(:,:,:,particleId))))/ ...
            (max(max(imageVec(:,:,:,particleId))) - min(min(imageVec(:,:,:,particleId))));
    catch
    end
end

% loading trained neural network
net = load(['neuralNetwork.mat']);
neuralNetwork = net.neuralNet;
classVec(:,1) = classify(neuralNetwork,imageVec,'ExecutionEnvironment','cpu','MiniBatchSize',100000,'Acceleration','auto');

% accepting or rejecting particles based on classification
trueParticlePosCount = 0;
for particleId = 1 : size(centroids,1)
    if classVec(particleId,1) == '1' && ...
            centroids(particleId,1) > 2 + roiRadius && ...
            centroids(particleId,1) < size(imageRaw,1) - roiRadius - 1 && ...
            centroids(particleId,2) > 2 + roiRadius && ...
            centroids(particleId,2) < size(imageRaw,2) - roiRadius - 1
        trueParticlePosCount = trueParticlePosCount + 1;
        app.data.file(fileId).particle(trueParticlePosCount).state = 'accepted';
        app.data.file(fileId).particle(trueParticlePosCount).centroid.x = centroids(particleId,1);
        app.data.file(fileId).particle(trueParticlePosCount).centroid.y = centroids(particleId,2);
    end
end

% merging particles
roiRadius = app.param.detection.roiRadius;
for particleId_1 = 1 : trueParticlePosCount
    if strcmp(app.data.file(fileId).particle(particleId_1).state,'accepted')
        for particleId_2 = 1 : trueParticlePosCount
            if strcmp(app.data.file(fileId).particle(particleId_2).state,'accepted') && particleId_1 ~= particleId_2
                if abs(app.data.file(fileId).particle(particleId_1).centroid.x - ...
                        app.data.file(fileId).particle(particleId_2).centroid.x) < ((2 * roiRadius) + 1) &&...
                        abs(app.data.file(fileId).particle(particleId_1).centroid.y - ...
                        app.data.file(fileId).particle(particleId_2).centroid.y) < ((2 * roiRadius) + 1)
                    app.data.file(fileId).particle(particleId_2).state = 'rejected';
                    break;
                end
            end
        end
    end
end
end

%%====================ParticleLocalizer=====================%%
function ParticleLocalizer(app,fileId)

% extracting number of particles
numParticles = length(app.data.file(fileId).particle);

% calling particleLocalizerAuxFn
if numParticles > 0
    particleLocalizerAuxFn(app,fileId);
end
end

%%====================ParticleRejector=====================%%
function ParticleRejector(app,fileId)

% extracting number of particles
numParticles = length(app.data.file(fileId).particle);

% calling particleRejectorAuxFn
if numParticles > 0
    particleRejectorAuxFn(app,fileId);
end
end

%%====================SpecificExport=====================%%
function SpecificExport(app)

% extracting ROI radius
roiRadius = app.param.detection.roiRadius;

% extracting number of files
numFiles = length(app.data.file);

for fileId = 1 : numFiles
    
    % creating new folder
    mkdir(fullfile(app.param.paths.calibrationAndUnknownData,app.data.file(fileId).type,'detected'));
    
    % extracting number of particles
    numParticles = length(app.data.file(fileId).particle);
    
    % extracting image
    image = uint16(app.data.file(fileId).image);
    
    % calculating ROI intensity
    roiInt = max(image(:));
    
    % looping through particles
    for particleId = 1 : numParticles
        
        if strcmp(app.data.file(fileId).particle(particleId).state,'accepted')
            
            % extracting particle centroid
            centroid.x = app.data.file(fileId).particle(particleId).centroid.x;
            centroid.y = app.data.file(fileId).particle(particleId).centroid.y;
            
            % assigning ROI pixels
            for x = round(centroid.x) - roiRadius : round(centroid.x) + roiRadius
                for y = round(centroid.y) - roiRadius : round(centroid.y) + roiRadius
                    if (x - round(centroid.x)) ^ 2 + (y - round(centroid.y)) ^ 2 > (roiRadius - 0.5) ^ 2 && ...
                            (x - round(centroid.x)) ^ 2 + (y - round(centroid.y)) ^ 2 < (roiRadius + 0.5) ^ 2
                        image(y,x) = roiInt;
                    end
                end
            end
        end
    end
    
    % exporting annotated images
    imwrite(image,...
        fullfile(app.param.paths.calibrationAndUnknownData,app.data.file(fileId).type,'detected',app.data.file(fileId).name),'WriteMode','overwrite');
    if fileId == 1
        imwrite(image,...
            fullfile(app.param.paths.calibrationAndUnknownData,app.data.file(fileId).type,'detected','all.tif'),'WriteMode','overwrite');
    else
        imwrite(image,...
            fullfile(app.param.paths.calibrationAndUnknownData,app.data.file(fileId).type,'detected','all.tif'),'WriteMode','append');
    end
end
end
```
