## Supplementary material for "Systematic assessment of the accuracy of subunit counting in biomolecular complexes using automated single molecule brightness analysis": exportFn.m

```
function returnFlag = exportFn(app)
% exportFn() -
% exports data generated for SAS.
%
% Syntax -
% exportFn(app,exportMode).
%
% Parameters -
% - app: SAS UI class
% - exportMode: SAS module name.

%% initializing returnFlag
returnFlag = false;

%% displaying SAS progress
app.msgBox.Value = sprintf('%s','Progress: export started.');

%% extracting number of files
numFiles = length(app.data.file);

%% writing files to disk
try
    
    %% looping through files
    for fileId = 1 : numFiles
        
        %% obtaining export folder path
        mkdir(fullfile(app.param.paths.calibrationAndUnknownData,app.data.file(fileId).type,'native'));
        
        %% setting up progress
        app.msgBox.Value = sprintf('%s',['Progress: exporting data in file ' num2str(fileId) ' out of ' num2str(numFiles) '.']);
        drawnow;
        
        %% saving .sd file
        dataArray = app.data.file(fileId);
        save([fullfile(app.param.paths.calibrationAndUnknownData,...
            app.data.file(fileId).type,...
            'native',...
            erase(app.data.file(fileId).name,{'.tif','.sd'})) '.sd'],'dataArray');
    end
catch
    returnFlag = true;
    app.msgBox.Value = sprintf('%s','Error: cannot write files to disk.');
    return
end
end
```
