## Supplementary material for "Systematic assessment of the accuracy of subunit counting in biomolecular complexes using automated single molecule brightness analysis": extractFn.m

```
function returnFlag = extractFn(app)
% extractFn() - 
% extracts data saved by SAS.
%
% Syntax - 
% extractFn(app).
%
% Parameters -
% - app: SAS UI class

%% initializing returnFlag
returnFlag = false;

%% initializing app properties
app.data = struct();

%% checking availability of sd files
inputFiles = dir(fullfile(app.param.paths.calibrationAndUnknownData,'**','*.sd'));
if isempty(inputFiles)
    returnFlag = true;
    app.msgBox.Value = sprintf('%s','Error: no sd files available.');
    return;
end

%% extracting number of files
numFiles = numel(inputFiles);

%% looping through files
for fileId = 1 : numFiles
    
    %% reading file name and folder
    if numFiles == 1
        fileName = inputFiles.name;
        fileFolder = inputFiles.folder;
    else
        fileName = inputFiles(fileId).name;
        fileFolder = inputFiles(fileId).folder;
    end
    filePath = fullfile(fileFolder,fileName);
    
    %% setting up SAS progress
    app.msgBox.Value = sprintf('%s',['Progress: extracting data from file ' num2str(fileId) ' out of ' num2str(numFiles) '.']);
    drawnow;
    
    %% replacing extension with .mat
    newFilePath = strrep(filePath,'.sd','.mat');
    
    %% copying file
    copyfile(filePath,newFilePath,'f');
    
    %% loading data
    try
        dataTemp = load(newFilePath);
    catch
        returnFlag = true;
        app.msgBox.Value = sprintf('%s',['Error: cannot load data from (' fileName ').']);
        return;
    end
    
    %% deleting copied file
    delete(newFilePath);
    
    %% assigning data array to app UI class particleData property
    app.data.file(fileId) = dataTemp.dataArray;
end

%% displaying SAS progress
app.msgBox.Value = sprintf('%s','Progress: extraction complete.');
drawnow;
end
```
