## Supplementary material for "Systematic assessment of the accuracy of subunit counting in biomolecular complexes using automated single molecule brightness analysis": main.m

```
function main(app)
% main entry point
% calls all SAS functions to detect, classify and analyze photobleaching traces
%
% Copyright 2020 John S H Danial
% Department of Chemistry, Univerity of Cambridge

% setting folder paths - EDITABLE
app.param.paths.calibrationAndUnknownData = app.CalibrationunknowndataEditField.Value;

% detection parameters - EDITABLE
app.param.detection.detect = app.DetectSwitch.Value == 'Y';
app.param.detection.cameraPixelSize = app.CameraPixelSizeEditField.Value;
app.param.detection.cameraOffset = app.CameraOffsetEditField.Value;
app.param.detection.cameraQE = app.CameraQEEditField.Value;
app.param.detection.cameraEMGain = app.CameraEMGainEditField.Value;
app.param.detection.maxSigma = app.MaximumSigmaEditField.Value;
app.param.detection.roiRadius = app.ROIRadiusEditField.Value;

% processing parameters - EDITABLE
app.param.processing.process = app.ProcessSwitch.Value == 'Y';

% annotation parameters - EDITABLE (only for calibration traces)
app.param.annotation.annotate = app.AnnotateSwitch.Value == 'Y';

% analysis parameters - EDITABLE
app.param.analysis.analyse = app.AnalyzeSwitch.Value == 'Y';
app.param.analysis.labelingEfficiency = app.LabelingeffeciencyEditField.Value;
app.param.analysis.maxGMM = app.MaximumGMMEditField.Value;
app.param.analysis.refine = app.RefineCheckBox.Value;

% setting returnFlag
returnFlag = false;

% running modules - DO NOT EDIT
if app.param.detection.detect && ~returnFlag
    returnFlag = detectFn(app);
end
if app.param.processing.process && ~returnFlag
    extractFn(app);
    returnFlag = processFn(app);
end
if app.param.annotation.annotate && ~returnFlag
    extractFn(app);
    returnFlag = annotateFn(app);
end

if app.param.analysis.analyse && ~returnFlag
    extractFn(app);
    analyzeFn(app);
end

% clearing RAM
app.data = struct();
end
```
