## Supplementary material for "Systematic assessment of the accuracy of subunit counting in biomolecular complexes using automated single molecule brightness analysis": processFn.m

```
function returnFlag = processFn(app)
% processFn() -
% extract traces of detected single molecules in field of view.
%
% Syntax -
% extractTraceFn(app).
%
% Parameters -
% - app: SAS UI class

%% initializing returnFlag
returnFlag = false;

%% extracting parameters
cameraOffset = app.param.detection.cameraOffset;
cameraEMGain = app.param.detection.cameraEMGain;
cameraQE = app.param.detection.cameraQE;

%% reading number of files
numFiles = length(app.data.file);

%% displaying SAS progress
app.msgBox.Value = sprintf('%s','Progress: extraction of traces started.');

%% looping through files
for fileId = 1 : numFiles
    
    %% reading file path
    if ~strcmp(app.data.file(fileId).state,'detected')
        returnFlag = true;
        app.msgBox.Value = sprintf('%s','Error: particles in found files were not detected.');
        return;
    end
    
    %% extracting file information and number of frames
    filePath = fullfile(app.param.paths.calibrationAndUnknownData,...
        app.data.file(fileId).type,...
        app.data.file(fileId).name);
    fileInfo = imfinfo(filePath);
    numFrames = numel(fileInfo);
    
    %% reading frame
    try
        video = zeros(size(imread(filePath,1),1),size(imread(filePath,1),2),numFrames);
        for frameId = 1 : numFrames
            video(:,:,frameId) = ((double(imread(filePath,frameId,'Info',fileInfo)) - cameraOffset) ./ cameraEMGain) ./ (cameraQE / 100);
        end
    catch
        returnFlag = true;
        app.msgBox.Value = sprintf('%s',['Error: cannot read image file (' app.data.file(fileId).name ').']);
        return;
    end
    
    %% setting up progress
    app.msgBox.Value = sprintf('%s',['Progress: processing particles in file ' num2str(fileId) ' out of ' num2str(numFiles)]);
    drawnow;
    
    %% calling ParticleDetector
    BrightnessMeasurer(app,fileId,video);
end

%% displaying progress
app.msgBox.Value = sprintf('%s','Progress: extraction of traces complete.');
drawnow;

%% exporting .sd files
exportFn(app);
end

%%====================BrightnessMeasurer=====================%%
function BrightnessMeasurer(app,fileId,video)

% extracting number of particles
numParticles = length(app.data.file(fileId).particle);

% extracting ROI radius
roiRadius = app.param.detection.roiRadius;

% looping through all files
for frameId = 1 : size(video,3)
    
    % looping through particles
    for particleId = 1 : numParticles
        
        % checking if particle is accepted
        if strcmp(app.data.file(fileId).particle(particleId).state,'accepted')
            
            % extracting centroid
            centroid.x = app.data.file(fileId).particle(particleId).centroid.x;
            centroid.y = app.data.file(fileId).particle(particleId).centroid.y;
            
            % extracting intensity and background
            app.data.file(fileId).particle(particleId).frame(frameId).intensity = sum(...
                video(round(centroid.y) - roiRadius : round(centroid.y) + roiRadius,...
                round(centroid.x) - roiRadius : round(centroid.x) + roiRadius,...
                frameId),'all');
            app.data.file(fileId).particle(particleId).frame(frameId).background = (sum(...
                video(round(centroid.y) - roiRadius - 2  : round(centroid.y) + roiRadius + 2,...
                round(centroid.x) - roiRadius - 2 : round(centroid.x) + roiRadius + 2,...
                frameId),'all') - ...
                app.data.file(fileId).particle(particleId).frame(frameId).intensity) ./...
                (((((roiRadius + 2) * 2) + 1) ^ 2) - (((roiRadius * 2) + 1) ^ 2)) .*...
                (((roiRadius * 2) + 1) ^ 2);
        else
            app.data.file(fileId).particle(particleId).frame(frameId).intensity = 0;
            app.data.file(fileId).particle(particleId).frame(frameId).background = 0;
            
        end
    end
end
for particleId = 1 : numParticles
    app.data.file(fileId).particle(particleId).maxIntensity = median([app.data.file(fileId).particle(particleId).frame(1:5).intensity]);
    app.data.file(fileId).particle(particleId).minIntensity = median([app.data.file(fileId).particle(particleId).frame(end - 4:end).background]);
end
end
```
