## Supplementary material for "Systematic assessment of the accuracy of subunit counting in biomolecular complexes using automated single molecule brightness analysis": SimulatedDataGenerator.m

```
%% Data simulator
%% Copyright 2020 John S H Danial
%% Department of Chemistry, Univerity of Cambridge

%% clears all variables and sets path - DO NOT EDIT
clear vars;
path = uigetdir;

%% camera parameters - EDITABLE
cameraPixelSize = 160;
cameraFrameSize = 256;
cameraEMGain = 300;
cameraOffset = 100;
cameraQE = 95;
cameraConversionFactor = 5.1;
cameraExposureTime = 0.03;
cameraDarkCurrent = 0.007 / (1 / cameraExposureTime);
cameraReadOut = 1;

%% sample parameters - EDITABLE
stoichiometry = [40 40];
numStructures = sum(stoichiometry);
photonCount = 10;
varPhotonCount = 0;
lateralPrecision = 130;

%% movie parameters - EDITABLE
numFrames = 500;
numMovies = 10;

%% looping through movies
for movieId = 1 : numMovies
    
    %% calling FrameGenerator
    [frameCoorX,frameCoorY,bleachTime,intensity] = FrameGenerator(numStructures,stoichiometry,cameraFrameSize,cameraPixelSize,photonCount,varPhotonCount,numFrames);
    
    %% calling StructureGenerator
    movie = MovieGenerator(frameCoorX,frameCoorY,bleachTime,intensity,lateralPrecision,cameraPixelSize,cameraFrameSize,cameraOffset,cameraQE,cameraEMGain,cameraConversionFactor,cameraDarkCurrent,cameraReadOut,numFrames);

    %% exporting movies
    for frameId = 1 : numFrames
        if frameId == 1
            imwrite(uint16(movie(:,:,frameId)),fullfile(path,[num2str(movieId) '.tif']),'WriteMode','overwrite');
        else
            imwrite(uint16(movie(:,:,frameId)),fullfile(path,[num2str(movieId) '.tif']),'WriteMode','append');
        end
    end
end

%%==========FrameGenerator===========%%
function [frameCoorX,frameCoorY,bleachTime,intensity] = FrameGenerator(numStructures,stoichiometry,cameraFrameSize,cameraPixelSize,photonCount,varPhotonCount,numFrames)

% initializing
globalCount = 1;

% calculating x and y coordinates of each structure
ranTransx = (((6 + 1) * cameraPixelSize) + ((cameraFrameSize - 6) * cameraPixelSize - ((6 + 1) * cameraPixelSize)) * rand(1,numStructures));
ranTransy = (((6 + 1) * cameraPixelSize) + ((cameraFrameSize - 6) * cameraPixelSize - ((6 + 1) * cameraPixelSize)) * rand(1,numStructures));

% calculating number of units in each structure
numUnits = [];
for unitId = 1 : length(stoichiometry)
    numUnits = [numUnits unitId .* ones(1,stoichiometry(unitId) * numStructures / sum(stoichiometry))];
end

% looping through structures
for structureId = 1 : numStructures
    
    % looping through units
    for unitId = 1 : numUnits(structureId)
        frameCoorX(globalCount) = ranTransx(structureId); 
        frameCoorY(globalCount) = ranTransy(structureId);
        globalCount = globalCount + 1;
    end
end

% creating randing bleach times and intensities
bleachTime = round(1 + (numFrames - 1) * rand(1,length(frameCoorX)));
intensity = normrnd(double(photonCount),varPhotonCount * double(photonCount),1,length(frameCoorX));
end

%%==========MovieGenerator===========%%
function movie = MovieGenerator(frameCoorX,frameCoorY,bleachTime,intensity,lateralPrecision,cameraPixelSize,cameraFrameSize,cameraOffset,cameraQE,cameraEMGain,cameraConversionFactor,cameraDarkCurrent,cameraReadOut,numFrames)

% creating kernel
for xVal = 1 : 11
    for yVal = 1 : 11
        kernel(yVal,xVal) = double(exp(-((xVal - 6) ^ 2 + (yVal - 6) ^ 2) /...
            (2 * ((lateralPrecision / cameraPixelSize) ^ 2))));
    end
end

% creating an empty movie array
movie(:,:,:) = zeros(cameraFrameSize,cameraFrameSize,numFrames);

% adding antibody and label length to frameCoor
currentframeCoorX = (frameCoorX ./ cameraPixelSize);
currentframeCoorY = (frameCoorY ./ cameraPixelSize);

% generating gaussian profiles
for unitId = 1 : size(frameCoorX,2)
    
    % adding kernel to movie
    xVal = round(currentframeCoorX(unitId)) - 5 : round(currentframeCoorX(unitId)) + 5;
    yVal = round(currentframeCoorY(unitId)) - 5 : round(currentframeCoorY(unitId)) + 5;
    movie(yVal,xVal,1:bleachTime(unitId)) = ...
        movie(yVal,xVal,1:bleachTime(unitId)) + (intensity(unitId) .* repmat(kernel,1,1,bleachTime(unitId)));
end

% converting photons to electrons
movie = (movie .* (cameraQE / 100)) + cameraDarkCurrent + cameraReadOut;

% adding noise and offset
movie = (gamrnd(movie,(cameraEMGain - 1 + (1 ./ movie))) ./ cameraConversionFactor) + cameraOffset;
end
```
