## Supplementary figures and images for "Systematic assessment of the accuracy of subunit counting in biomolecular complexes using automated single molecule brightness analysis"

### splash.png

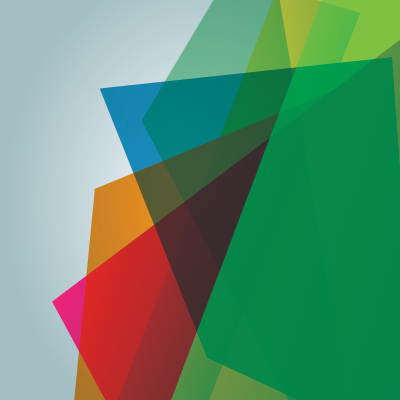
